# Variable precision in visual perception

**DOI:** 10.1101/153650

**Authors:** Shan Shen, Wei Ji Ma

## Abstract

Given the same sensory stimuli in the same task, human observers do not always make the same response. Well-known sources of behavioral variability are sensory noise and guessing. Visual short-term memory studies have suggested that the precision of the sensory noise is itself variable. However, it is unknown whether precision is also variable in perceptual tasks without a memory component. We searched for evidence for variable precision in 11 visual perception tasks with a single relevant feature, orientation. We specifically examined the effect of distractor stimuli: distractors were absent, homogeneous and fixed across trials, homogeneous and variable, or heterogeneous and variable. We first considered four models: with and without guessing, and with and without variability in precision. We quantified the importance of both factors using six metrics: factor knock-in difference, factor knock-out difference, and log factor posterior ratio, each based on AIC or BIC. According to all six metrics, we found strong evidence for variable precision in five experiments. Next, we extended our model space to include potential confounding factors: the oblique effect and decision noise. This left strong evidence for variable precision in only one experiment, in which distractors were homogeneous but variable. Finally, when we considered suboptimal decision rules, the evidence also disappeared in this experiment. Our results provide little evidence for variable precision overall and only a hint when distractors are variable. Methodologically, the results underline the importance of including multiple factors in factorial model comparison: testing for only two factors would have yielded an incorrect conclusion.

## INTRODUCTION

When presented with the same stimuli in the same perceptual task, human observers do not always make the same response. One source of such variability is noise in the encoding stage – the mapping from the stimulus to the internal representation. This mapping is noisy at the neural level (Faisal, Selen, & Wolpert, 2008; London, Roth, Beeren, Häusser, & Latham, 2010; Tolhurst, Movshon, & Dean, 1983) and has long been modeled as noisy in behavioral models (Fechner, 1860; Green & Swets, 1966; Thurstone, 1927). It is furthermore common to assume that such sensory or encoding noise follows a Gaussian distribution in the stimulus space (Green & Swets, 1966), or a Von Mises distribution when the stimulus variable is circular (Wilken & Ma, 2004; Zhang & Luck, 2008).

In recent years, the idea has been explored that encoding precision – roughly the inverse of the variance of the sensory noise – is itself a random variable. Such random variability is distinct from the systematic variation of precision with set size (Mazyar, van den Berg, & Ma, 2012; Mazyar, van den Berg, & Seilheimer, 2013; Palmer, 1990; Shaw, 1980; Wilken & Ma, 2004); throughout this paper, variability in precision will refer to variability at a given set size. Variable-precision models have been used to model human (Donkin, Nosofsky, Gold, & Shiffrin, 2013; Fougnie, Suchow, & Alvarez, 2012; keshvari, van den Berg, & Ma, 2012, 2013; Oberauer & Lin, 2017; Pratte, Park, Rademaker, & Tong, 2017; van den Berg, Awh, & Ma, 2014; van den Berg, Shin, Chou, George, & Ma, 2012) and monkey (D. T. Devkar, Wright, & Ma, 2015; D. Devkar, Wright, & Ma, 2017) behavior in visual short-term memory tasks as well as human behavior in visual search tasks (Bhardwaj, van den Berg, Ma, & Josić, 2016; Mazyar et al., 2012, 2013). A related concept appears in the beta-binomial model for the psychometric curve (Schütt, Harmeling, Macke, & Wichmann, 2016), where an extra parameter is used to capture variability in the probability of a binary response. At the neural level, variable precision might have a parallel in the single (Bays, 2014) or double stochasticity of neural spike counts (Churchland et al., 2011; Goris, Movshon, & Simoncelli, 2014; van den Berg, Yoo, & Ma, 2017).

Variable precision could in principle be confounded with or partly explained by other factors. First, on some proportion of trials, encoding precision might be exactly zero, for example due to lapses in attention; this is typically modeled as a guessing rate (Harvey, 1986; Watson & Pelli, 1983; Wichmann & Hill, 2001). Moreover, in binary decisions, errors in the mapping between decision and motor output are mathematically equivalent to guesses. Since variable precision in a sense interpolates between zero precision and a fixed non-zero precision, it sometimes mimics guessing (keshvari et al., 2013; van den Berg et al., 2014, 2012). Second, variability in precision could be partly explained by systematic variations of precision across the stimulus space. For example, cardinal orientations (horizontal or vertical) are encoded with higher precision than oblique orientations. This phenomenon is called the “oblique effect” (Andrews, 1965, 1967; Appelle, 1972; Girshick, Landy, & Simoncelli, 2011; Pratte et al., 2017) and is an example of *heteroskedasticity*, whereby some measure of dispersion (skedasis) differs across subgroups. Heteroskedasticity has also been described in color perception and color visual short-term memory (Bae, Olkkonen, Allred, Wilson, & Flombaum, 2014). Heteroskedasticity could be due to a non-uniform distribution of the preferred stimuli of visual cortical neurons (Li et al., 2003; De Valois, William Yund, & Hepler, 1982; Mansfield & Ronner, 1978). This distribution might in turn have adapted to stimulus statistics in natural environments (Attneave, 1954; Barlow, 1961; Girshick et al., 2011) and can be explained by theories of efficient coding (Ganguli & Simoncelli, 2014; Wei & Stocker, 2015). Third, decision noise or suboptimality in inference might be confounded with sensory noise in general, and with variable precision in particular. Decision noise refers to any noise in the mapping from the internal representation to the decision (Mueller & Weidemann, 2008). Statistical inefficiency or inference noise (Burgess, Wagner, Jennings, & Barlow, 1981; Drugowitsch, Wyart, Devauchelle, & koechlin, 2016; Liu, knill, & kersten, 1995), model mismatch (Beck, Ma, Pitkow, Latham, & Pouget, 2012; Orhan & Jacobs, 2014), and other forms of systematic suboptimal inference (Gigerenzer & Goldstein, 1996; Shen & Ma, 2016; Simon, 1956) could mimic decision noise, because a model that models the decision stage as optimal will attribute any systematic deviations from optimality to random variability, i.e. decision noise.

It is in principle possible that variability in precision found in previous work captures what is in reality guessing, heteroskedasticity, or decision noise. Only a few studies, all in the realm of visual short-term memory, have attempted to disentangle some of these factors. Some have compared a variable-precision model to a fixed-precision model with a lapse rate (D. T. Devkar et al., 2015; keshvari et al., 2012; van den Berg et al., 2014, 2012). Others have argued that the oblique effect accounts for most of what otherwise would be designated as variable precision (Pratte et al., 2017). Here, we attempt to distinguish guessing, the oblique effect, and decision noise from residual variable precision by including all factors simultaneously in our models.

## EXPERIMENTAL METHODS

### Experimental design

We conducted eight new target categorization experiments and analyzed the results of three previously published experiments (**Table 1**, **Figure 1**). The previously published experiments are numbered Experiment 7 (was Experiment 1 in Shen & Ma, 2016), Experiment 8 (was Experiment 2 in Mazyar et al., 2013), and Experiment 11 (was Experiment 1 in Mazyar et al., 2013). All experiments were identical in the following aspects:

- Stimuli were Gabors, with orientation the only relevant feature.
- Subjects fixated and all stimuli were presented at the same eccentricity (5° of visual angle).
- Stimuli were presented for a short duration (50 or 83 ms).
- There was no substantial visual short-term memory component.
- Subjects made binary choices.
- There were no intertrial dependencies.

**Figure 1.**
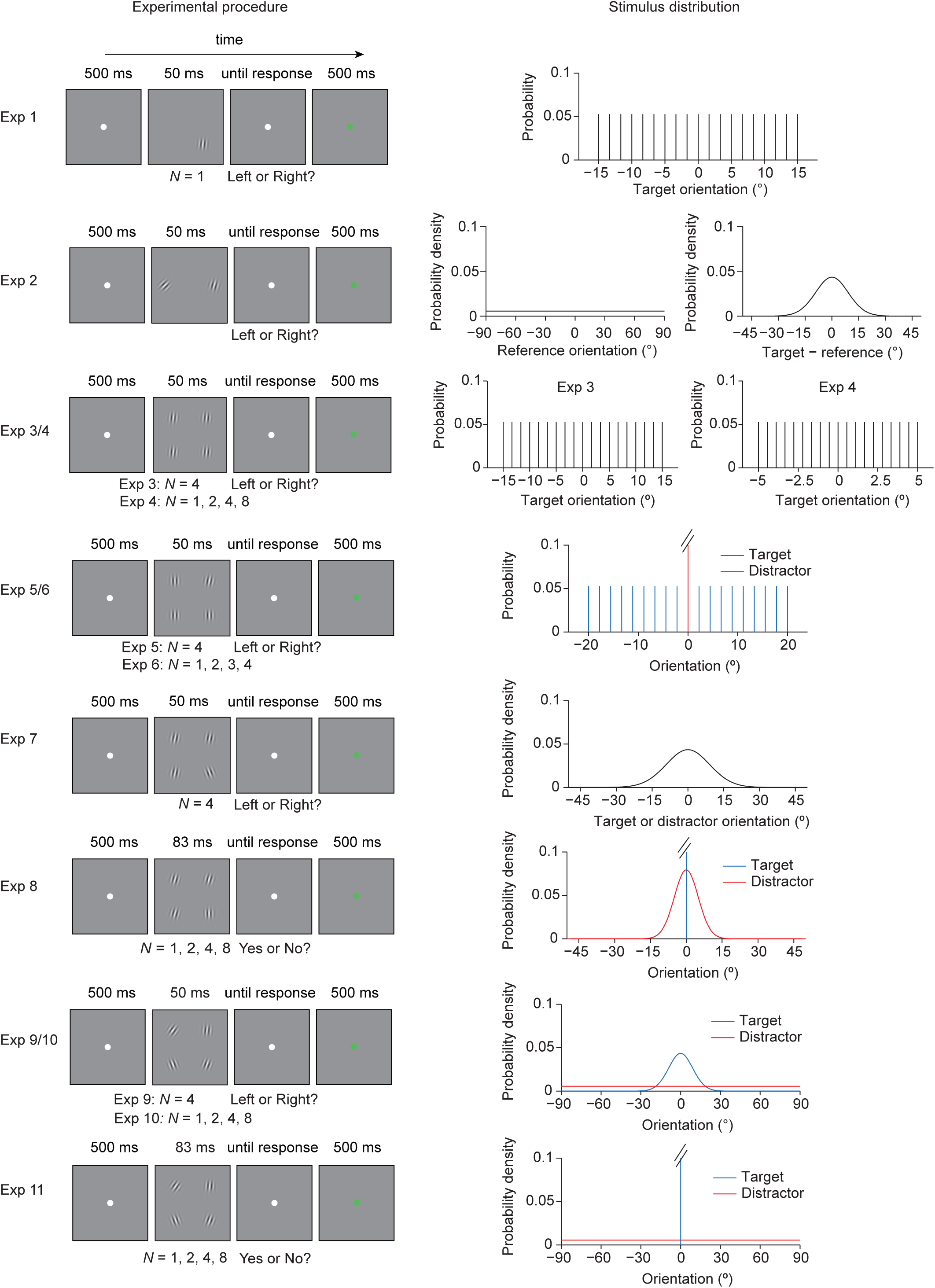
Experimental designs. The left column shows the trial procedure and the right column shows the orientation distribution of the stimuli.

We designed our experiments to search for variability in precision that cannot be accounted for by set size, guessing, the oblique effect, or decision noise. In the realm of visual short-term memory, it has been suggested that precision is variable due to stochasticity in the rate of decay of memory (Fougnie et al., 2012); however, our study is not a memory study. Another idea has been that stimulus context has a large effect on the quality of any one stimulus (Brady & Alvarez, 2015). We take inspiration from this suggestion and examine, whether evidence for residual variable precision is stronger in experiments where the stimulus context is more variable. Concretely, we considered a variety of visual perception tasks that differed in the complexity of the distractor context (**Table 1**): in Experiments 1 to 4, there were no distractors; in Experiments 5 and 6, distractors were homogeneous and their value remained unchanged over trials; in Experiments 7 and 8, distractors were homogeneous but varied across trials; and finally, in Experiments 9 to 11, distractors were both heterogeneous and variable across trials. We hypothesized that the evidence for the “residual” variable precision is higher when the distractor context is more complex.

Besides in distractor context, our experiments differed in other aspects, including task type (detection or categorization), orientation range (narrow or full), number of targets (1 or multiple), set size (single or multiple stimuli), and set size context (single or multiple set sizes). To the best of our ability, we examined the effects of these factors, to ensure that our conclusions are robust.

## Apparatus and stimuli

Subjects were seated at a viewing distance of approximately 60 cm. All stimuli were displayed on a 21-inch LCD monitor with a refresh rate of 60 Hz and a resolution of 1280×1024 pixels. The stimulus displays were composed of Gabor patches shown on a grey background. In Experiments 1-7, 9, and 10, background luminance was 29.3 cd/m^2^ and the Gabors had a peak luminance of 35.2 cd/m^2^, a spatial frequency of 3.1 cycles per degree of visual angle, a standard deviation of the Gaussian envelope of 0.25° of visual angle, and a phase of 0 for the cosine pattern. Settings were different in Experiments 8 and 11 (see Mazyar et al., (2013)): background luminance was 33.1 cd/m^2^ and the Gabors had a peak luminance of 122 cd/m^2^, a spatial frequency of 1.6 cycles per degree of visual angle, a standard deviation of the Gaussian envelope of 0.29° of visual angle, and a phase of 0 for the cosine pattern.

## Experimental procedure

Each trial started with a fixation dot on a blank screen (500 ms), followed by a stimulus display (50 ms in Experiments 1-7, 9, and 10; 83 ms in Experiments 8 and 11). Then, a blank screen was shown until the subject responded by pressing a button. Response time was not limited. After the response, correctness feedback was given by changing the color of the fixation dot (green for correct, red for incorrect; 500 ms; **Figure 1**). Experiments 1-7, 9, and 10 were visual categorization tasks and Experiment 8 and 11 were visual detection tasks.

Experiments 1-7, 9, and 10 each consisted of three sessions on different days. Each session consisted of five blocks, and each block contained 200 trials, for a total of 3×5×200=3000 trials per subject. All blocks were statistically identically to each other.

Experiment 8 (Experiment 2 in Mazyar et al. (2013)) consisted of four sessions; here, we analyze only the two sessions with homogeneous distractors (for details, see Mazyar et al. (2013)). Each session consisted of four blocks, and each block contained 175 trials, for a total of 1400 trials per subject analyzed here.

Experiment 11 (Experiment 1 in Mazyar et al. (2013)) consisted of six sessions; here, we analyze only the two sessions with “High” heterogeneity (for details, see Mazyar et al. (2013)). Each session consisted of four blocks, and each block contained 175 trials, for a total of 1400 trials per subject analyzed here.

## Stimulus displays and tasks

We now describe the stimulus display in each of the 11 experiments (**Figure 1**). We will use the phrase “drawn randomly” as shorthand for “drawn randomly from a uniform distribution over the values specified”. The radial positions of all stimuli in all experiments were 5° of visual angle relative to fixation. For the angular positions of the stimuli, we use the standard convention of polar coordinates: 0° corresponds to the positive horizontal axis, and positive values correspond to positions counterclockwise with respect to that axis. For stimulus orientations, we use the convention that is most natural given our orientation distributions: 0° is vertical and positive values are clockwise.

*Experiment 1*: The stimulus display consisted of a single stimulus in one of four angular positions: −135°, −45°, 45°, and 135°. Stimulus orientation was drawn randomly from 19 values equally spaced between −15° and 15°. The subject reported the tilt with respect to vertical of a single oriented stimulus.

*Experiment 2*: The stimulus display consisted of two stimuli, placed on the horizontal axis to the left and right of the fixation. The stimulus on the right was the reference stimulus, whose orientation *s*_ref_ was drawn randomly (from a uniform distribution over the entire orientation space). The stimulus on the left was the target stimulus, whose orientation was drawn randomly from a Von Mises distribution centered at *s*_ref_ with a concentration parameter of 10. The subject reported whether the target was oriented clockwise or counterclockwise with respect to *s*_ref_.

*Experiments 3 and 4*: All stimuli were targets, and the subject reported the tilt of their common orientation. In Experiment 3, set size was 4, and the angular positions were as in Experiment 1. The common orientation was drawn randomly from 19 values equally spaced between −15° and 15°. In Experiment 4, set size was 1, 2, 4, or 8, drawn randomly. Angular positions were chosen as follows, in order to maximize spacing. At set size 8, we used all 8 angular positions: 0°, 45°, 90°, 135°, 180°, −45°, −90°, −135°. At set sizes 1, 2, and 4, we placed the first stimulus at a random angular position. At set size 2, we then placed the second stimulus diametrically opposite to the first, while at set size 4, we placed the remaining 3 stimuli at every other position. Stimulus orientation was drawn randomly from 19 values equally spaced between −5° and 5°.

*Experiment 5 and 6*: Experiment 5 and 6 were target classification tasks. Subjects reported the tilt of the target relative to vertical. In Experiment 5, set size was 4 and the angular positions were the same as in Experiment 1. Three of the stimuli were vertical; these were the distractors. The fourth stimulus, whose position was drawn randomly from the four positions, was the target. Target orientation was drawn randomly from 19 values equally spaced between −20° and 20°. The design of Experiment 6 was identical to that of Experiment 5, except that set size was 1, 2, 3, or 4, drawn randomly. Angular positions were drawn randomly.

*Experiment 7:* We reanalyzed data from a previously published target classification experiment (Shen & Ma, 2016). Set size was 4 and the angular positions were the same as in Experiment 1. Each display contained one target and three distractors; target position was drawn randomly from the four positions. The target orientation and the common distractor orientation were drawn independently from the same Von Mises distribution centered at vertical, with a concentration parameter of 10 (similar to a Gaussian distribution with a standard deviation of 9.06°). Subjects reported the tilt of the target (the unique stimulus).

*Experiment 8:* Experiment 8 was an orientation detection task. Subjects reported whether or not a target was present. Set size was 1, 2, 4, or 8, drawn pseudorandomly. At set size 8, all angular positions were used. At set sizes 1, 2, and 4, the first stimulus was placed at a random angular position, and the remaining stimuli were placed at adjacent positions. The target orientation was vertical. Trial type was “target present” or “target absent”, drawn pseudorandomly. On target-absent trials, all stimuli were distractors. On target-present trials, one stimulus was the target stimulus and the remaining stimuli were distractors; the position of the target stimulus was drawn randomly from the available positions. The common orientation of the distractors was drawn from a Von Mises distribution centered at vertical, with a concentration parameter of 32 (similar to a Gaussian distribution with a standard deviation of 5.06°).

*Experiment 9 and 10*: Experiments 9 and 10 were target classification tasks. Subjects reported the tilt of the target relative to vertical. In Experiment 9, set size was 4 and the angular positions were the same as in Experiment 1. Each stimulus display contained one target and three distractors; target position was drawn randomly. Target orientation was drawn from a Von Mises distribution with a mean of 0 and a concentration parameter of 10 (similar to a Gaussian distribution with a standard deviation of 9.06°). Each of the distractor orientations was drawn independently from a uniform distribution over the entire orientation space. The tasks in these experiments contained ambiguity, in the sense that on some trials, either answer could be correct even in the absence of sensory noise, because of the overlap between the target and distractor distributions; as experimenters, we set the tilt of the generated target as the correct answer. To help subjects learn the task, we provided 10 static example Gabor patches from the target and distractor distributions and verbally explained that the distractors were more likely to have large tilts than the targets. The subjects performed well above chance (71.7 ± 1.6%, Student’s *t*-test: *t*(5) = 13.9, *p* < 10^−4^). The design of Experiment 10 was identical to Experiment 9, except that the set size was 1, 2, 4, or 8, drawn randomly on each trial. The stimulus placement was the same as in Experiment 4. This experiment combined distractors that were variable both within and across trials with multiple set sizes. Again, this experiment had ambiguity when the set size was greater than 1, and therefore did not allow for perfect performance. Nevertheless, subject accuracy was 72.6 ± 1.7%, well above chance (Student’s *t*-test: *t*(10) = 13.1, *p* < 10^−6^).

*Experiment 11*: We reanalyzed data from a previously published target detection task (Experiment 1 in Mazyar et al., 2013). The basic paradigm was the same as in Experiment 8, except that the distractors were heterogeneous. Each distractor was independently drawn from a uniform distribution over the entire orientation space, which was the same as in Experiments 9 and 10. This experiment differed from those not only in the type of task (detection instead of categorization), but also in the absence of ambiguity: the stimulus statistics did not preclude perfect performance.

**Table 1:** Overview of experiments. For distractors, we use “homogeneous” and “heterogeneous” to indicate that the distractors are identical to (or different from, respectively) each other within a display; we use “variable” to indicate variability across trials. Experiment 7 was previously published as Experiment 1 in Shen & Ma (2016). Experiments 8 and 11 were previously published as Experiments 2 and 1 in Mazyar et al. (2013), respectively.

| Experiment | Number of subjects | Number of stimuli | Number of targets | Task | Distractors |
| --- | --- | --- | --- | --- | --- |
| 1 | 6 | 1 | 1 | Target categorization relative to vertical | None |
| 2 | 5 | 2 | 1 | Target categorization relative to reference (stimulus on the right) | None |
| 3 | 6 | 4 | all | Target categorization relative to vertical | None |
| 4 | 6 | 1, 2, 4, 8 | all | Target categorization relative to vertical | None |
| 5 | 6 | 4 | 1 | Target categorization relative to vertical | Vertical |
| 6 | 6 | 1, 2, 3, 4 | 1 | Target categorization relative to vertical | Vertical |
| 7 | 10 | 4 | 1 | Target categorization relative to vertical | Homogeneous, variable |
| 8 | 13 | 1, 2, 4, 8 | 0, 1 | Target detection, vertical target | Homogeneous, variable |
| 9 | 6 | 4 | 1 | Target categorization relative to vertical | Heterogeneous, variable |
| 10 | 11 | 1, 2, 4, 8 | 1 | Target categorization relative to vertical | Heterogeneous, variable |
| 11 | 6 | 1, 2, 4, 8 | 0, 1 | Target detection,<br>vertical target | Heterogeneous,<br>variable |

## THEORY

For each experiment, we build process models in which the stimuli give rise to measurements and the observer applies a decision rule to the measurements to produce a category estimate (**Figure 2A**). Our models consist of three steps:

1. The generative model: a statistical description of the noisy internal measurements of the stimuli and of the observer’s beliefs about how the stimuli are generated (which may or may not be how they are actually generated). In this step, we consider two kinds of variability in the precision of the measurement noise: the oblique effect (O) and “residual” variable precision (V).
2. The observer’s decision model. In each experiment, we assume that the observer applies an optimal decision rule, which maximizes the posterior distribution under the generative model of Step 1. In some experiments, we also consider alternative suboptimal decision rules. Note that even the optimal decision rule is not optimal in an absolute sense, because (a) measurement noise is present, (b) the generative model assumed by the observer is not necessarily identical to the true generative model, (c) we allow for the decision to be corrupted by decision noise (D).
3. Prediction for the probabilities of the possible subject responses on a given trial, i.e. given the stimulus values on that trial). This combines the stimulus-conditioned measurement distributions from Step 1 with the decision rule from Step 2. We also incorporate guessing (G) in this step.

**Figure 2.**
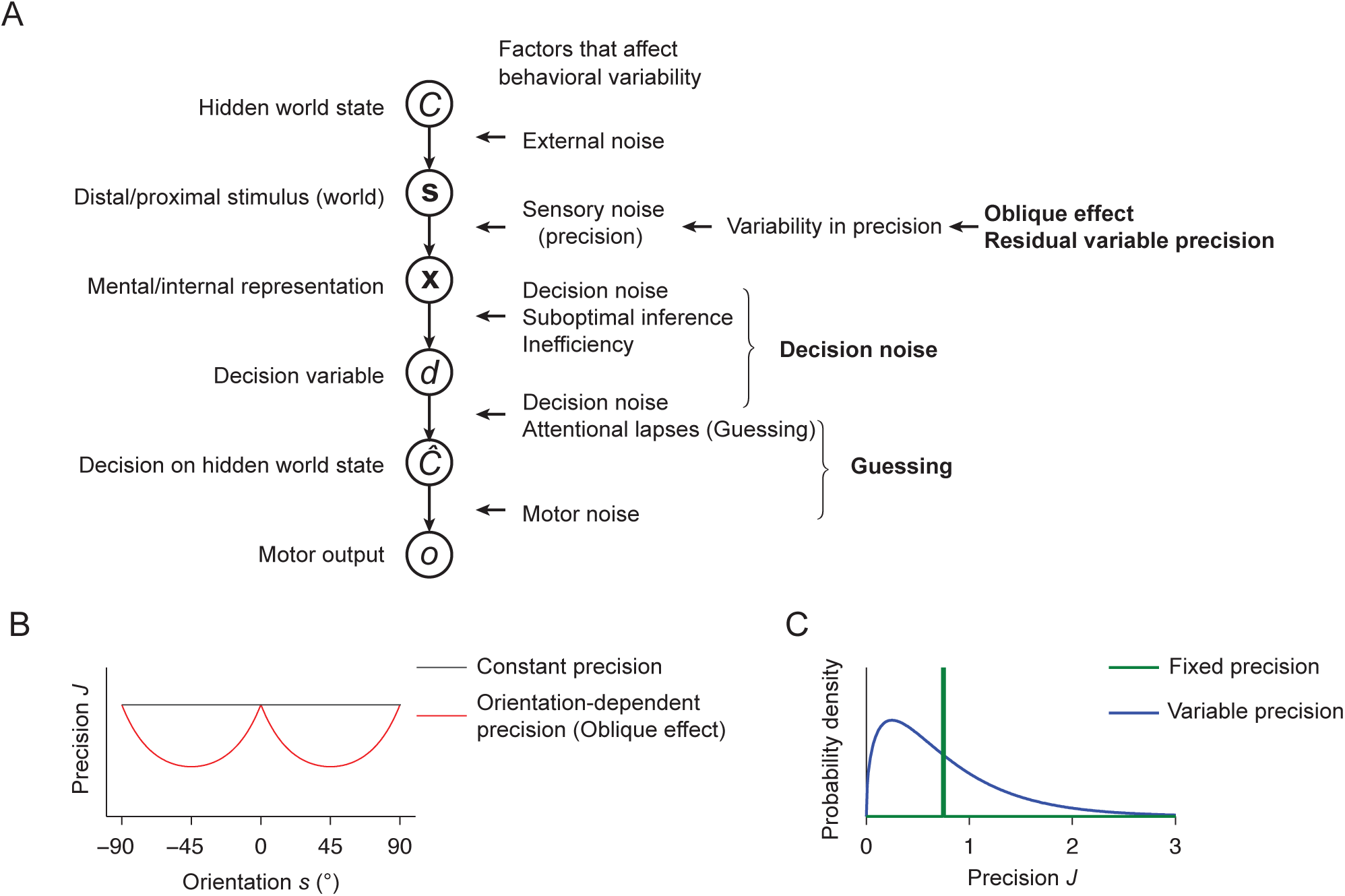
Generative model and factors that might affect behavioral variability. **(A)** The diagram shows the generic generative model of our tasks. Each node represents a variable and each arrow between two nodes represents a conditional dependence. Factors that might affect behavioral variability are listed to the right of the diagram. Here, we test the bold-faced ones: oblique effect, residual variable precision, decision noise and guessing. **(B)** We model the dependence of precision *J* on orientation *s* (the oblique effect) as 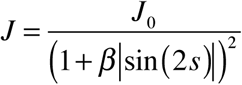 (red). The black line represents constant precision (*β*=0). **(C)** In variable-precision models, we model the probability distribution over precision as a gamma distribution; an example with a mean of 0.75 and a scale parameter *τ* of 0.5 is shown in blue. The green line represents a delta function over precision, corresponding to fixed precision (*τ* =0).

We now describe each of these steps in greater detail.

### Step 1a: Generative model: Noisy measurements

We assume that the observer makes a noisy measurement *x_i_* of each physical orientation *s_i_*, where *i* =1,…, *N* labels the stimuli in a given display (*N* is the set size). We denote the vector of physical orientations of the stimuli by **s** and the vector of orientation measurements by **x**. Throughout, we assume that the measurements are independent given the stimuli,

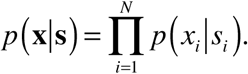

Depending on the stimulus range (narrow or wide), we assume that the distribution of *x*_*i*_ given *s_i_* is either Gaussian,

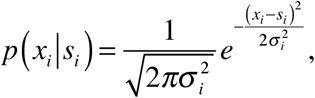

or Von Mises (circular Gaussian),

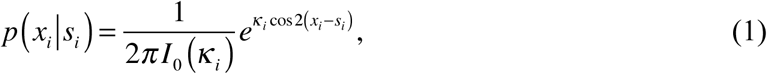

where *I*_0_ is the modified Bessel function of the first kind of order 0. Noise level or precision is controlled by the standard deviation σ_*i*_ (Gaussian) or by the concentration parameter *κ_i_* (Von Mises). The factor 2 in the exponent of the Von Mises distribution appears because orientation space is [0, π) instead of [0, 2π). In the limit of large *κ_i_* the Von Mises distribution converges to the Gaussian distribution, with 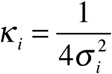. We use the Gaussian distribution when the range of orientations (experiments other than Experiment 2) or the orientation difference (Experiment 2) in the experiment is small compared to the full orientation space of [0, π) (in Experiments 1 to 8), and Von Mises otherwise (in Experiments 9 to 11).

A general definition of precision based on *p*(*x*_*i*_|*s*_*i*_) is as Fisher information (Cover & Thomas, 2005). Fisher information, denoted by *J*, is related to the parameters above through

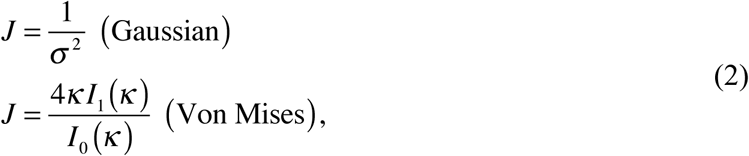

where *I*_1_ is the modified Bessel function of the first kind of order 1. In previous work (keshvari et al., 2012; Mazyar et al., 2012; Mazyar et al., 2013; van den Berg et al., 2012), we did not include the factor of 2 in **Eq. (1)** and the factor of 4 in **Eq. (2)**, but instead rescaled orientations from [0, π) to [0, 2π) before doing any analysis. This rescaling is mathematically equivalent to inserting those factors, but here, we opted against the rescaling, so that we can compare the results of Gaussian-based analysis to those of Von Mises-based analysis with minimal confusion (e.g. in **Figure 13**).

### Step 1b: Generative model: Variability in precision

Next, we consider variability in the precision of the measurement noise. This variability can be due to multiple sources.

*Oblique effect (O)*. To model the oblique effect, we introduce stimulus dependence in the dispersion parameter of the measurement distribution. For Gaussian noise, we take the standard deviation of the noise to be a rectified sine function of the stimulus orientation (Girshick et al., 2011):

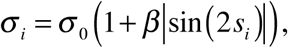

where σ_0_ is the baseline noise level and β is the amplitude parameter of orientation dependence. When β=0, there is no oblique effect. For precision *J_i_*, we obtain (**Figure 2B**):

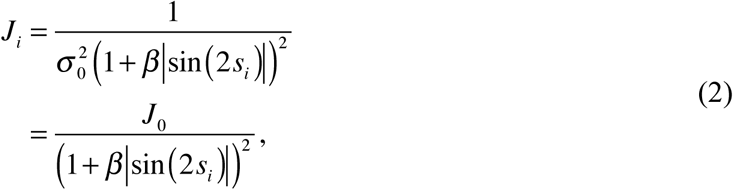

where 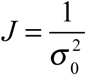 is the baseline precision. We use the latter equation also for Von Mises noise.

*“Residual” variable precision (V)*. Besides the oblique effect, precision might vary for other reasons; we will consider all other sources collectively and call them “residual” variable precision, denoted by V. Variable-precision models have been successful in describing behavior in many visual short-term memory tasks, including delayed estimation (Fougnie et al., 2012; van den Berg et al., 2014, 2012), change detection (keshvari et al., 2012, 2013), and change localization (D. T. Devkar et al., 2015; keshvari et al., 2012). Most of these papers have formalized variability in precision by a assuming a gamma distribution over *J:*

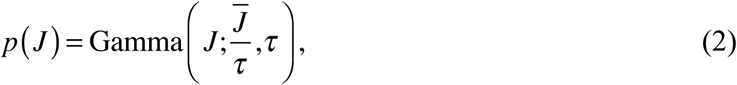

where 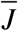 is the mean precision, and τ is called the scale parameter (**Figure 2C**). We will follow this formalism here.

*Combining factors O and V*. In all experiments, we tested all four combinations of the two forms of precision variability: a base model with fixed precision (Base), a model with only the oblique effect (O), a model with only residual variable precision (V), and a model with both (OV). In the Base model, *J_i_* is the same across stimuli *i* and across trials. In the O model, *J_i_* is computed from **Eq. (2)**. In the V model, *J_i_* is drawn independently across *i* and across trials from a gamma distribution with mean 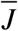 and scale parameter τ (**Eq. (2)**). In the OV model, we first compute 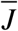 from **Eq. (2)**, then draw *J* from a gamma distribution with mean 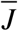 and scale parameter τ.

In experiments with multiple set sizes, we allowed *J* (models with fixed precision), *J*_0_ (models with the oblique effect), or 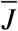 (models with residual variable precision) to vary with set size; we did not impose a parametric form but fitted the parameter independently at each set size (Mazyar et al., 2012, 2013).

### Step 1b: Generative model: Experimental statistics

The generative model consists not only of the distribution *p*(*x_i_*|*s_i_*), but also of the observer’s beliefs about the experimental statistics. The variables relevant to those beliefs are category (target tilted left or right in the categorization experiments, target present or absent in the detection experiments), and the individual stimulus orientations. We assume that the observer’s beliefs about the category distribution and the category-conditioned stimulus distributions are identical to the true ones, i.e. the ones set by the experimental design, with two exceptions:

- The two categories were always presented with probability 0.5. However, we did not assume that subjects would believe this probability to be exactly 0.5. Instead, we used a free parameter to characterizing the observer’s prior probability that the stimulus was tilted right (*p*_right_) in the categorization experiments, or that the stimulus was present (*p*_present_) in the detection experiments.
- In Experiments 1, 3, 4, 5, 6, we used discrete stimulus values, e.g. 19 values spaced linearly between −15° and 15° (Experiments 1 and 3), between −5° and 5° (Experiment 4), between −20° and 20° (Experiments 5 and 6). We did not assume that subjects had detailed knowledge of these values, but we instead assumed that the observer believed that this distribution was Gaussian with the same mean and standard deviation as the actual distribution. In Results, we examine this assumption.

### Step 2: Decision model: Optimal observer and decision noise

The Bayesian observer “inverts” the generative model to obtain a probability distribution over the variable of interest (here category, *C*=1 or *C*=-1 given the noisy measurements **x** on a given trial. The Bayesian decision variable, denoted by *d*, is the log of the ratios of the probabilities of *C*=1 and *C*=-1 given **x**:

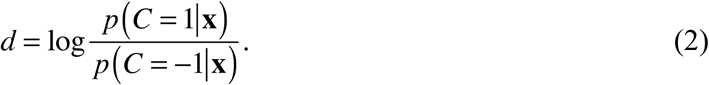

The Bayesian observer without decision noise reports *C*=1 if *d* is positive. The Bayesian observer is not strictly optimal, because we made two modifications in Step 1b; for a detailed distinction between the terms Bayesian and optimal, see Ma (2012).

The derivations of the Bayesian decision rules for all experiments are given in Appendix 1. In models with the oblique effect (O), we assume that the observer knows the noise level *σ* when evaluating **Eq. (2)**, but does not “realize” that there is a relationship between *σ* and orientation, *s*. Therefore, the observer does not infer *s* from *σ*, for example by marginalizing over *s*. For a more principled examination of the implications of heteroskedasticity for Bayesian observer models, see Wei & Stocker (2015).

*Decision noise (D)*. Decision noise (D) has been modeled using a softmax function (Daw, O’Doherty, Dayan, Seymour, & Dolan, 2006; Soltani, 2006), as Gaussian noise on the decision criterion (Mueller & Weidemann, 2008), or as Gaussian noise on the log posterior ratio (Drugowitsch et al., 2016; keshvari et al., 2012, 2013). Here, we use the last approach: the decision variable 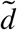 follows a Gaussian distribution with a mean of *d* (**Eq. (2)**) and a standard deviation of *σ*_d_:

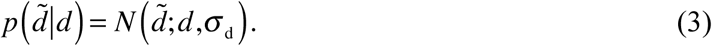

The observer reports *C*=1 if 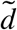 is positive.

### Step 3: Predictions: Sampling of measurements and guessing rate

Step 2 produces a mapping from a set of measurements, **x**, to an estimate of category Ĉ. However, we are ultimately interested in the probability that on a given trial, the observer will make either category response, that is, *p*(Ĉ|**s**), where **s** are the physical stimuli on that trial. This distribution is obtained as an average (marginalization) over measurement vectors **x**:

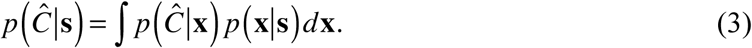

Here, *p*(Ĉ|**x**) is deterministic and given by Step 2, and *p*(**x**|**s**) is given by the measurement distributions in Step 1a. To approximate this integral, we sampled, for each trial in the experiment, a large number of measurement vectors **x** based on the physical stimuli **s** on that trial. For each **x**, we applied the decision rule from Step 2, and counted the outcomes. The proportions of either category response serve as our approximation of *p*(Ĉ|**s**). The number of samples of **x** needs to be sufficiently large for the approximation to be good. Based on an earlier test that showed convergence near 256 samples in a similar task (van den Berg et al., 2012, Appendix), we chose 2000 samples.

*Guessing (G)*. We allowed for the possibility that the subject guesses on some proportion of trials. To this end, we introduced a guessing rate λ, so that the probability of reporting Ĉ given **s** becomes

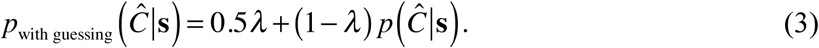

0.5 comes from the assumption that guesses are equally distributed across the responses.

*Factorial model comparison*. We will denote the factors by G, O, D and V (see **Table 2**). We tested these factors in a factorial manner (Acerbi, Vijayakumar, & Wolpert, 2014; van den Berg et al., 2014). We will denote each model by the combinations of factors in the model. For example, GDV has all factors except for O. This ended up models of 16 combinations. In some experiments, we will combine these combinations with both optimal versus suboptimal decision rules, but in most experiments, we will only consider the optimal decision rule.

**Table 2:**
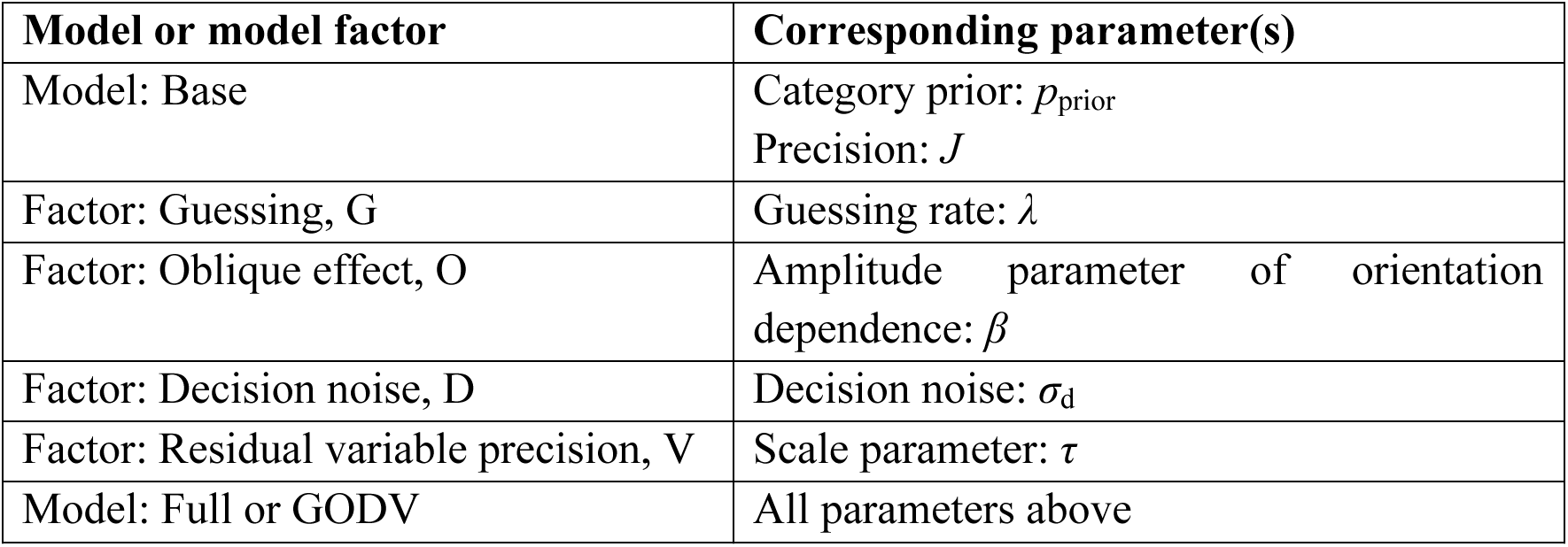
Models, model factors, and parameters. The Base model has parameters *J* and *p*_prior_. G, O, D, and V denote the model factors that can be added to the Base model, each with an associated parameter. The Full or GODV model is obtained by adding all four factors. In each model, we fitted all parameters on an individual-subject basis.

| Model or model factor | Corresponding parameter(s) |
| --- | --- |
| Model: Base | Category prior: $p_{\text{prior}}$<br>Precision: $J$ |
| Factor: Guessing, G | Guessing rate: $\lambda$ |
| Factor: Oblique effect, O | Amplitude parameter of orientation dependence: $\beta$ |
| Factor: Decision noise, D | Decision noise: $\sigma_d$ |
| Factor: Residual variable precision, V | Scale parameter: $\tau$ |
| Model: Full or GODV | All parameters above |

## MODELING METHODS

### Model fitting

We fitted the free parameters in each model (**Table 2**) to each individual subject’s data using maximum-likelihood estimation. The log likelihood of a given parameter combination is the logarithm of the probability of all of the subject’s responses given the model and each parameter combination:

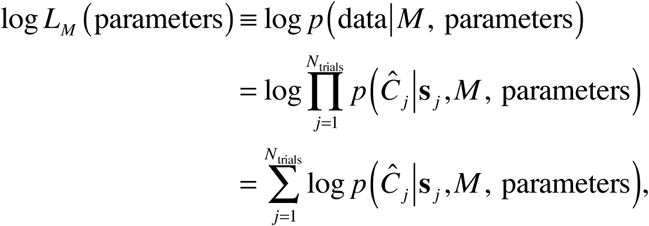

where *j* is the trial index, *N*_trials_ is the number of trials, **s**_*j*_ is the set of orientations presented on the *j* trial, *C_j_* is the subject s response on the *j*^*th*^ trial, and we have assumed that there are no sequential dependencies between trials. The probability of the subject response, log*p*(Ĉ_*j*_|**s**_j_, *M*, parameters), is obtained from **Eqs. (3)** or **(3)**. To find the values of parameters that maximize log *L_M_*(parameters), we used Bayesian Adaptive Direct Search (BADS, Acerbi & Ma, 2017), initialized with random values for all parameters. After BADS returned a parameter combination, we recomputed the log likelihood 10 times with that combination and took the mean, to reduce sampling noise. We performed this process for 10 different initializations and took the maximum of the log likelihoods as the maximum log likelihood for the model, LL_max_(*M*).

### Model comparison metrics

We use the Akaike Information Criterion (AIC, (Akaike, 1974)) and the Bayesian Information Criterion (BIC, (Schwarz, 1978)) as metrics of badness of fit. These metrics penalize a model for having more free parameters:

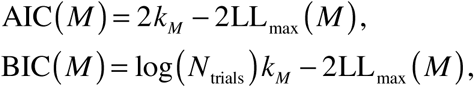

where *k_M_* is the number of parameters of model *M* and *N*_trials_ is the number of trials. Both AIC and BIC have their own advantages and disadvantages (Burnham & Anderson, 2002, pp. 293-305, for a pro-AIC account; kass & Raftery, 1995, for a pro-BIC account). AIC penalizes each parameter by 2 points, while BIC penalizes each parameter by 8.0 points in Experiments 1-7, 9, and 10, and by 7.2 points in Experiment 8 and 11. We only draw conclusions when both metrics provide strong evidence (see section “Jeffreys’ scale”). We use AIC/BIC and the corresponding factor importance metrics (below) for all formal conclusions. For some models in some experiments, we also show fits to the psychometric curves, but these are only meant as qualitative visual checks of the relative and absolute goodness of fit of the models.

### Factor importance metrics

We consider four model factors: guessing (G), the oblique effect (O), decision noise (D), and “residual” variable precision (V). Each can have two levels (absent and present), for a total of 16 models. We would like to draw conclusions about the importance of each factor regardless of model. We are also interested in the combination of O and V, because both are forms of variable precision; in the context of factor importance metrics, we will for brevity also refer to this combination as a factor.

In van den Berg et al. (2014), we quantified factor importance by calculating the proportion of subjects for whom all models in a given “model family” (e.g. all models in which G is absent) are rejected (according to AIC), as a function of the rejection criterion. This method has two disadvantages: a) it works at the population level and cannot be applied when the number of subjects is small; b) it outputs a curve (function) rather than a number. Therefore, we introduce three new factor importance metrics here (**Figure 3**): knock-in difference (representing evidence for factor usefulness), knock-out difference (representing evidence for factor necessity), and log factor likelihood ratio (representing evidence for factor presence). The terms “useful” and “necessary” only refer to goodness of fit, not to usefulness or necessity to the observer.

**Figure 3.**
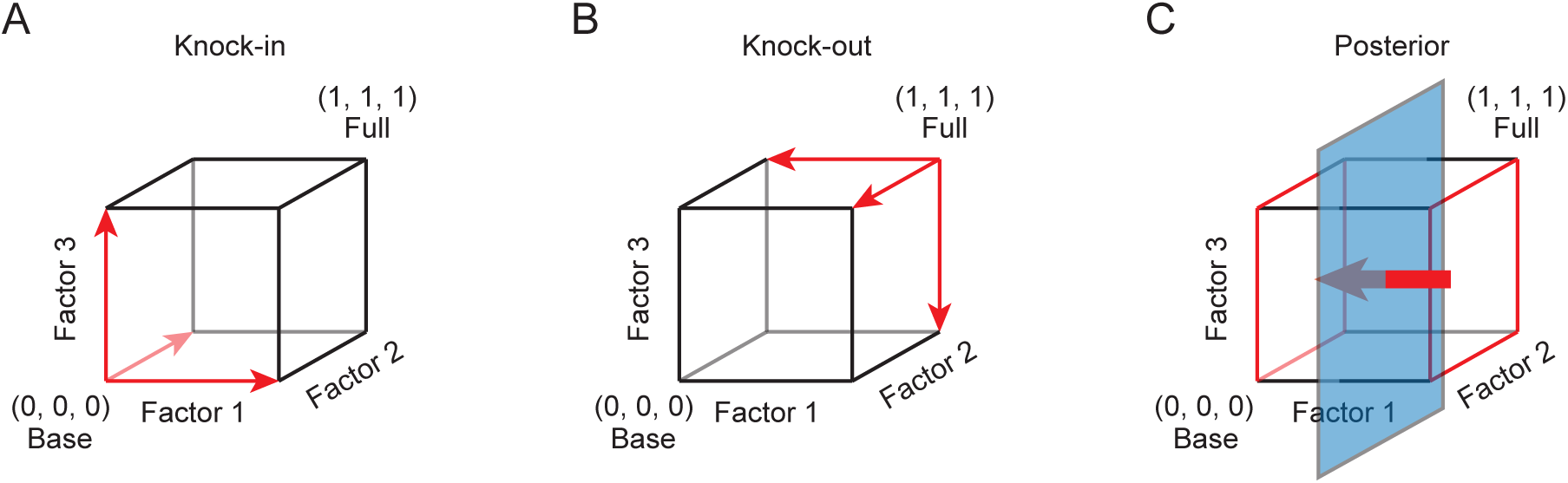
Factor importance metrics. In each diagram, each dimension represents a binary factor and each vertex a model; we show an example with 3 factors and thus a total of 8 models. The Base model, with none of the factors, is (0, 0, 0) and the Full model, with all factors, is (1, 1, 1). **(A)** knock-in difference (KID, red arrows): the AIC or BIC difference between the Base model (0, 0, 0) and the knock-in model with each single factor. **(B)** knock-out difference (KOD, red arrows): the AIC or BIC difference between the corresponding knock-out model and the Full model (1, 1, 1). **(C)** The log factor likelihood ratio (LFLR). We compute the log likelihood ratio of a factor being present versus absent by marginalizing over all models with or without that factor, respectively.

#### Factor usefulness: knock-in difference (KID)

We measure the evidence that a factor is useful as the amount by which the goodness of fit improves relative to the Base model by adding, or “knocking in”, that factor (**Figure 3A**). We define the *knock-in difference based on AIC* (KID_AIC_) of a factor *F* (which takes values G, O, D, V, or OV) as the AIC difference between the Base model and the knock-in model with *F*, denoted by “Base + *F*”:

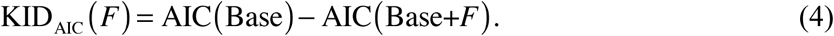

A positive KID_AIC_ means that the knock-in model fits better than the Base model, and represents evidence that the factor is useful. Drugowitsch et al. (2016) applied a similar analysis.

We call KID a measure of the “usefulness” of a factor and not of its “sufficiency”, because we take “insufficiency” of a model to refer to the deviation between the model and the true distribution (as estimated using deviance (Wichmann & Hill, 2001) or kullback-Leibler divergence (Shen & Ma, 2016)), which is not what we quantify here.

#### Factor necessity: knock-out difference (KOD)

We measure the evidence that a factor is necessary as the amount by which the goodness of fit of the Full model (GODV) decreases by lesioning, or “knocking out”, that factor (**Figure 3B**). We define the *knock-out difference based on AIC* (KOD_AIC_) of a factor *F* as the AIC difference between the corresponding knock-out model (ODV, GOV, GDV, GOD, or GD, denoted by “Full-*F*”) and the Full model:

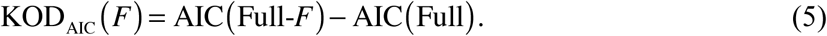

A positive KOD_AIC_ means that the knock-out model fits worse than the Full model, and represents evidence that the factor is necessary.

#### Factor presence: log factor likelihood ratio (LFLR)

Finally, we estimate the evidence that a factor is present in the true model underlying a subject’s behavior as the log likelihood ratio of a factor being present versus absent, which we will refer to as the log factor likelihood ratio (LFLR; **Figure 3C**). Although this quantity reflects most objectively the degree of belief in a factor (Van Horn, 2003), additional assumptions are needed to estimate it. To find the marginal likelihood that a factor *F* is present, we marginalize over all models *M* in the model space:

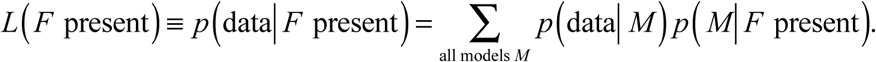

Next, we assume that the models we tested are representative, so that the sample average is a good approximation of the theoretical average:

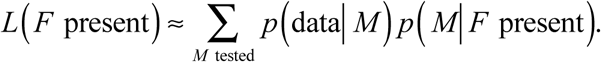

Next, we assume that all models containing *F* are a priori equally probable, so that

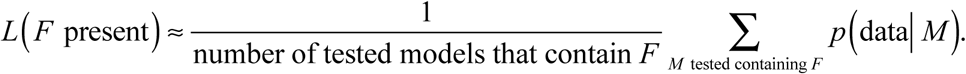

Finally, we approximate the log marginal likelihood of a given model by −0.5 times the AIC of that model (Akaike, 1978, 1979; Burnham & Anderson, 2002, Chapter 2.9):

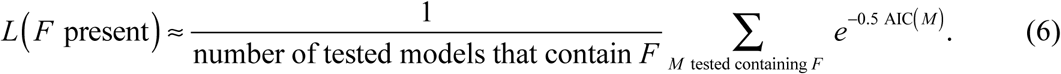

We analogously define the marginal likelihood of factor absence, and then the *log factor likelihood ratio based on AIC* (LFLR_AIC_) as

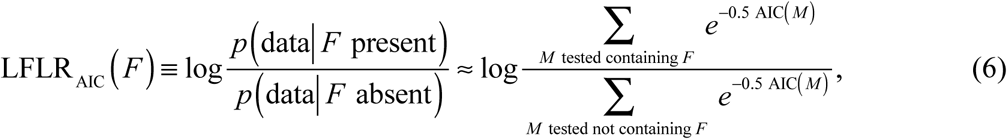

where we have assumed that equal numbers of tested models contain and do not contain *F*, as is the case throughout this paper. LFLR_AIC_ is similar to the log evidence ratio of AIC weights (E.-J. Wagenmakers & Farrell, 2004), except that the latter is an estimate of the log likelihood ratio between two models, instead of between factor presence and absence.

We now discuss an important special case. If adding a factor does not improve the unpenalized goodness of fit, which means that the model containing the factor has the exact same LL_max_ as the corresponding model without that factor, then its LFLR_AIC_ is:

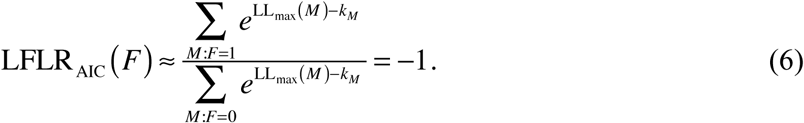

Therefore, the LFLR_AIC_ of a factor should always be higher than −1, but in practice, it is possible to be slightly lower because of the simulation noise.

Similarly, we could also use −0.5 BIC as an approximation of log marginal likelihood. Then, **Eqs. (4)**, **(5)**, **(6)** and **(6)** would be analogous, and the lower bound of LFLR_BIC_ in **Eq. (6)** would be −4.0 (Experiments 1-7, 9 and 10), or −3.6 (Experiments 8 and 11), dependent on the number of trials. In practice, BIC penalizes extra parameters by more than AIC. Therefore, LFLR_BIC_ is generally lower than LFLR_AIC_ for the same factor.

To facilitate comparison with KID and KOD, we will report the value of 2-LFLR rather than LFLR itself. For each metric, a higher positive value means more evidence that the factor is important.

#### Relation between KID and KOD

While the KID and KOD metrics each have a relatively straightforward interpretation, the question arises what inconsistency between them could mean. Finding that a factor is important in KID but not in KOD could indicate a “trade-off between factors, or a logical OR operation: the effect of knocking out one factor is compensated for by other factors, to yield an equally good fit. Such a “trade-off between factors is an example of model mimicry (Townsend, 1972; E. J. Wagenmakers, Ratcliff, Gomez, & Iverson, 2004) and would go away in the limit of infinite data. The opposite is also possible: a factor is important in KOD, but not in the KID. This could indicate an “interaction” between factors or a logical AND operation: neither factor is useful by itself, but their combination is, similar to finding an interaction without main effects in ANOVA.

#### Relation between LFLR and KID/KOD

In general, we expect LFLR to be more closely related to KOD than to KID. This is because the log-sum-exponent operation in the calculation of LFLR, **Eq. (6)**, is similar to a max operation (Ma, Shen, Dziugaite, & van den Berg, 2015). Thus, the marginal likelihoods of factor presence and absence will often be dominated by the best models with and without the factor, respectively. Take LFLR_AIC_ as an example. Starting from **Eq. (6)**,

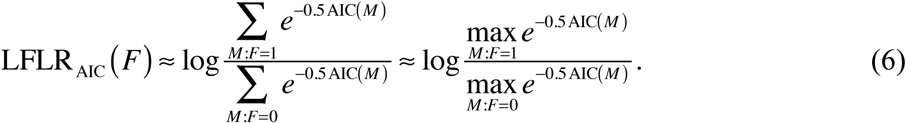

If furthermore, the lowest-AIC model is the most highly parameterized model, which is the most likely case, then 2-LFLR_AIC_ becomes identical to KOD_AIC_

#### Jeffreys’ scale

To interpret the numerical values of the factor importance metrics, we use Jeffreys’ scale (Jeffreys, 1961; **Table 3a**), which is commonly used to interpret Bayes factors. We make a few modifications (**Table 3b**): a) to be conservative, we are more careful with our adjectives than Jeffreys; b) also to be conservative, we base our interpretation on the lowest of FIM_AIC_ and FIM_BIC_; c) our scale is not symmetric between positive and negative values, because FIM has a lower bound due to being based on AIC or BIC (see **Eq. (6)**).

**Table 3a:** Jeffreys’ scale for Bayes’ factors (Jeffreys, 1961).

| Bayes Factor (BF) | 2·log(BF) | Interpretation |
| --- | --- | --- |
| >100 | >9.2 | Extreme evidence for H1 |
| 30 to 100 | 6.8 to 9.2 | Very strong evidence for H1 |
| 10 to 30 | 4.6 to 6.8 | Strong evidence for H1 |
| 3 to 10 | 2.2 to 4.6 | Moderate evidence for H1 |
| 1 to 3 | 0 to 2.2 | Anecdotal evidence for H1 |
| 1 | 0 | No evidence |
| 1/3 to 1 | -2.2 to 0 | Anecdotal evidence for H0 |
| 1/10 to 1/3 | -4.6 to -2.2 | Moderate evidence for H0 |
| 1/30 to 1/10 | -6.8 to -4.6 | Strong evidence for H0 |
| 1/100 to 1/10 | -9.2 to -6.8 | Very strong evidence for H0 |
| <1/100 | <-9.2 | Extreme evidence for H0 |

**Table 3b:** Jeffreys’ scale adapted for our three factor importance metrics (FIMs). “Important” can mean “useful” (KID), “necessary” (KOD), or “present” (LFLR).

| $\min(\text{FIM}_{\text{AIC}}(F), \text{FIM}_{\text{BIC}}(F))$ | Interpretation |
| --- | --- |
| >9.2 | Very strong evidence that $F$ is important |
| 6.8 to 9.2 | Strong evidence that $F$ is important |
| 4.6 to 6.8 | Moderate evidence that $F$ is important |
| <4.6 (including <0) | Little or no evidence that $F$ is important |

## RESULTS

Motivated by the visual short-term memory literature, we searched for evidence for variable precision in 11 experiments, most of which used visual search tasks. In doing so, we tested for three factors that could be confounded with variability in precision, namely guessing (G), the oblique effect (O), decision noise (D), and in some cases, suboptimal decision rules. Our approach relies on quantitative model comparison, the results of which we summarize through three novel “factor importance metrics”, each crossed with AIC and BIC.

### The importance of variable precision (V) when taking into account guessing (G)

Guessing, representing stimulus-independent lapses of attention or motor errors, is a factor that has been widely accepted to be present in psychophysical tasks, and it is routinely included in psychometric curve fits (Wichmann & Hill, 2001). In the current study, we started searching evidence for variable precision by only considering G. This ended up with four models: base model with no factors (Base), variable precision model (V), fixed precision with guessing (G), variable precision with guessing (GV). We applied all six factor importance metrics to this model set.

In most experiments (Experiments 1, 3, 5, 6, 7, 9, 10, and 11), mean KID_AIC_(V) and mean KID_BIC_(V) were both greater than 9.2 (**Figure 4A**), indicating very strong evidence that factor V is useful to explain the data. This is consistent with the model fits to the psychometric curves (**Figure 4D**, **Figures A1**, **A3**, **A5**, **A6**, **A7**, **A9**, **A10**, **panel B**, compare the Base and V models) In Experiments 2 and 4, KID_AIC_(V) and KID_BIC_(V) were much smaller, indicating little or no evidence that factor V is useful.

**Figure 4.**
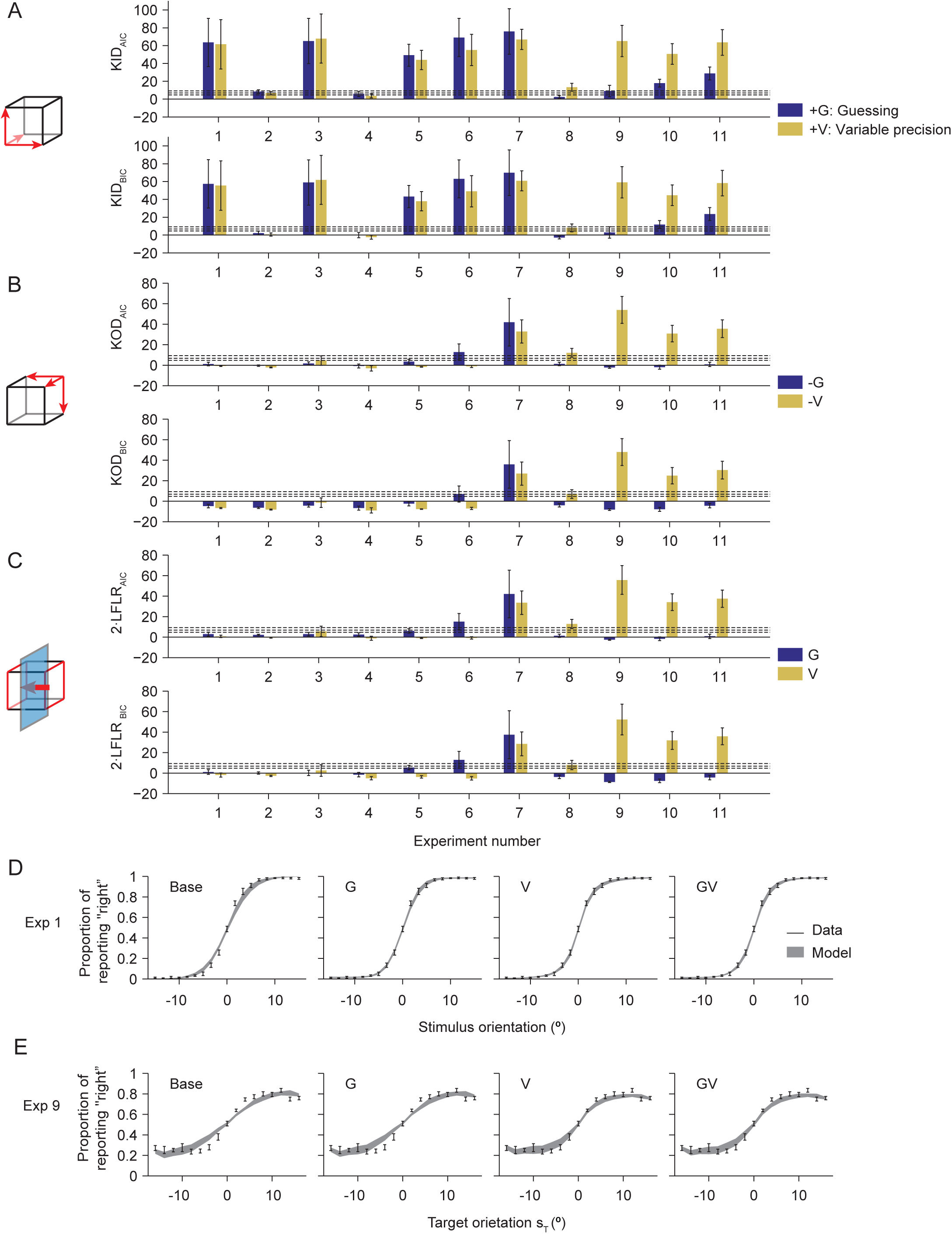
Factor importance: guessing (G) and variable precision (V). Here and in other factor importance plots, dashed black lines mark the boundaries of our interpretation of the strength of the evidence (>9.2: very strong, >6.8: strong, >4.6: moderate). **(A-C)** Mean and s.e.m. of KID (A), KOD (B), and 2·LFLR (C) based on AIC (top) or BIC (bottom) for the factors G and V in all experiments. **(D)** Model fits to the proportion of reporting “right” as a function of target orientation in Experiment 1. In all model fit plots, we use error bars and shaded areas to represent ± 1 s.e.m. in the data and the model fits, respectively. The G, V, and GV models fit the data equally well, and better than the Base model. **(E)** Model fits in Experiment 9. The V and GV models fit the data almost equally well, and better than the Base and G models.

**Figure A1.**
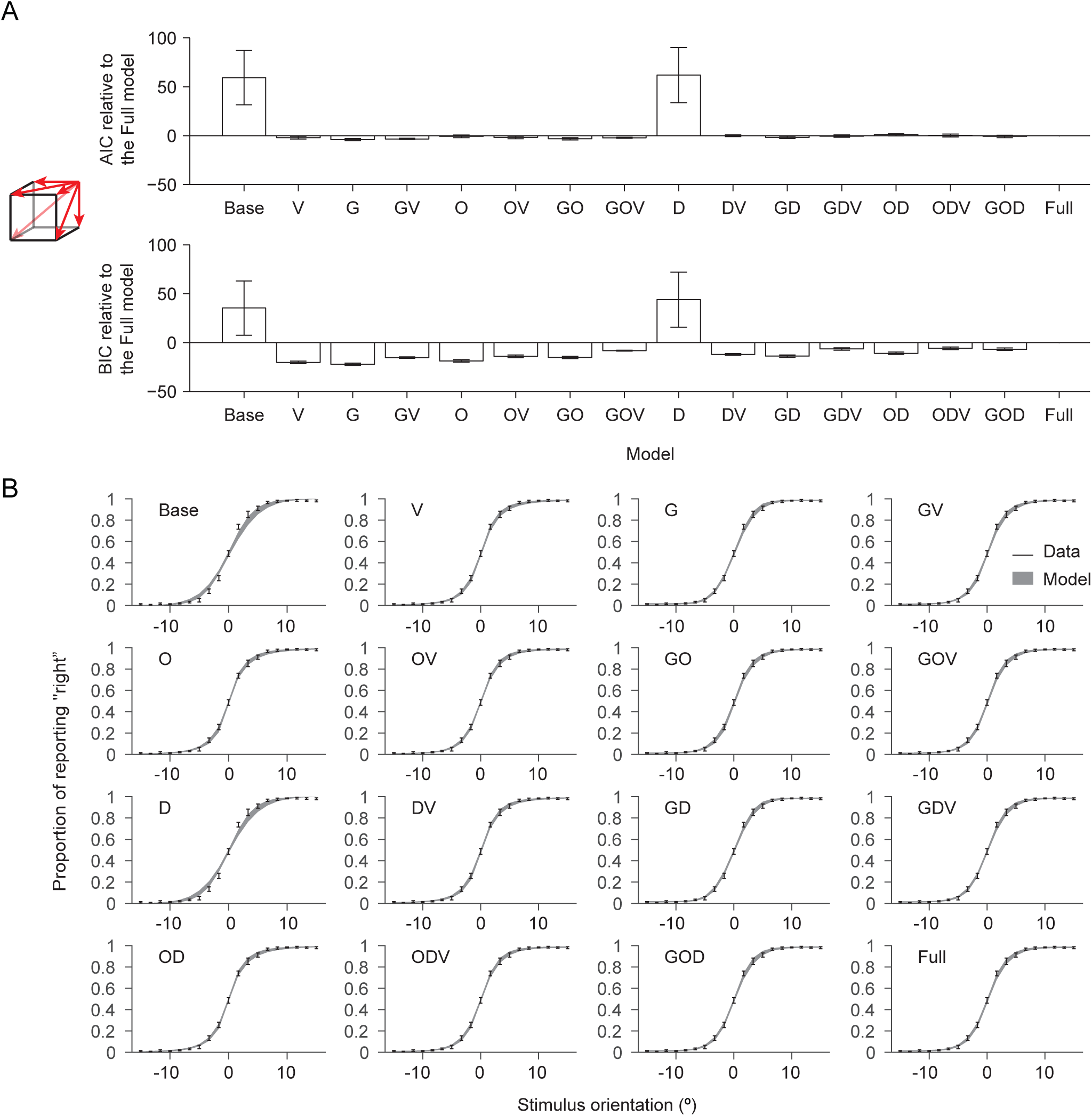
Experiment 1. **(A)** Complete model comparison. Mean and s.e.m. of the difference in AIC (top) and BIC (bottom) between each model and the Full model. **(B)** Proportion of reporting “right” as a function of stimulus orientation: data and model fits.

**Figure A2.**
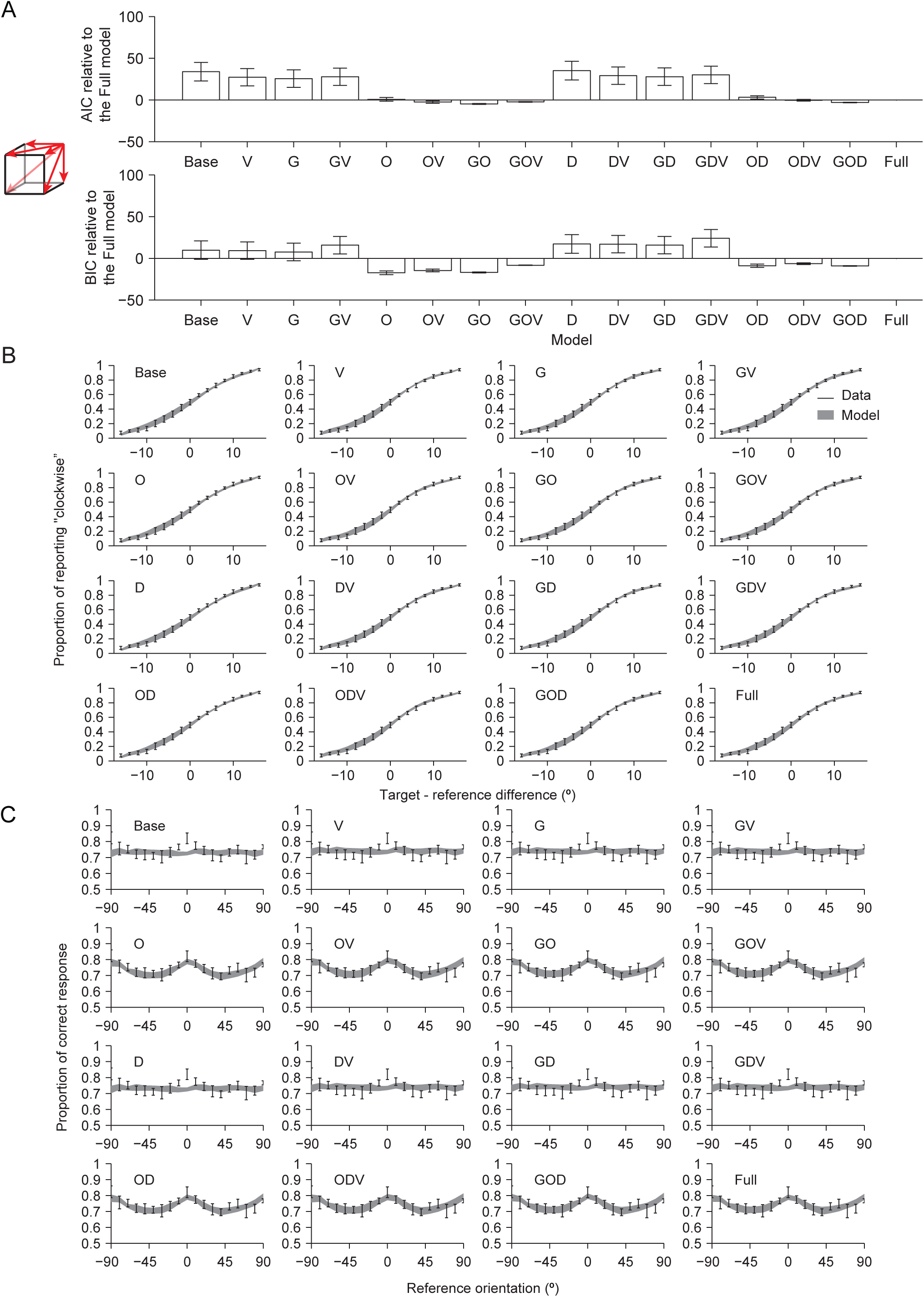
Experiment 2. **(A)** Complete model comparison. Mean and s.e.m. of the difference in AIC (top) and BIC (bottom) between each model and the Full model. **(B)** Proportion of reporting “clockwise” as a function of orientation difference between the target and the reference. **(C)** Proportion correct as a function of the reference orientation: data and model fits.

**Figure A3.**
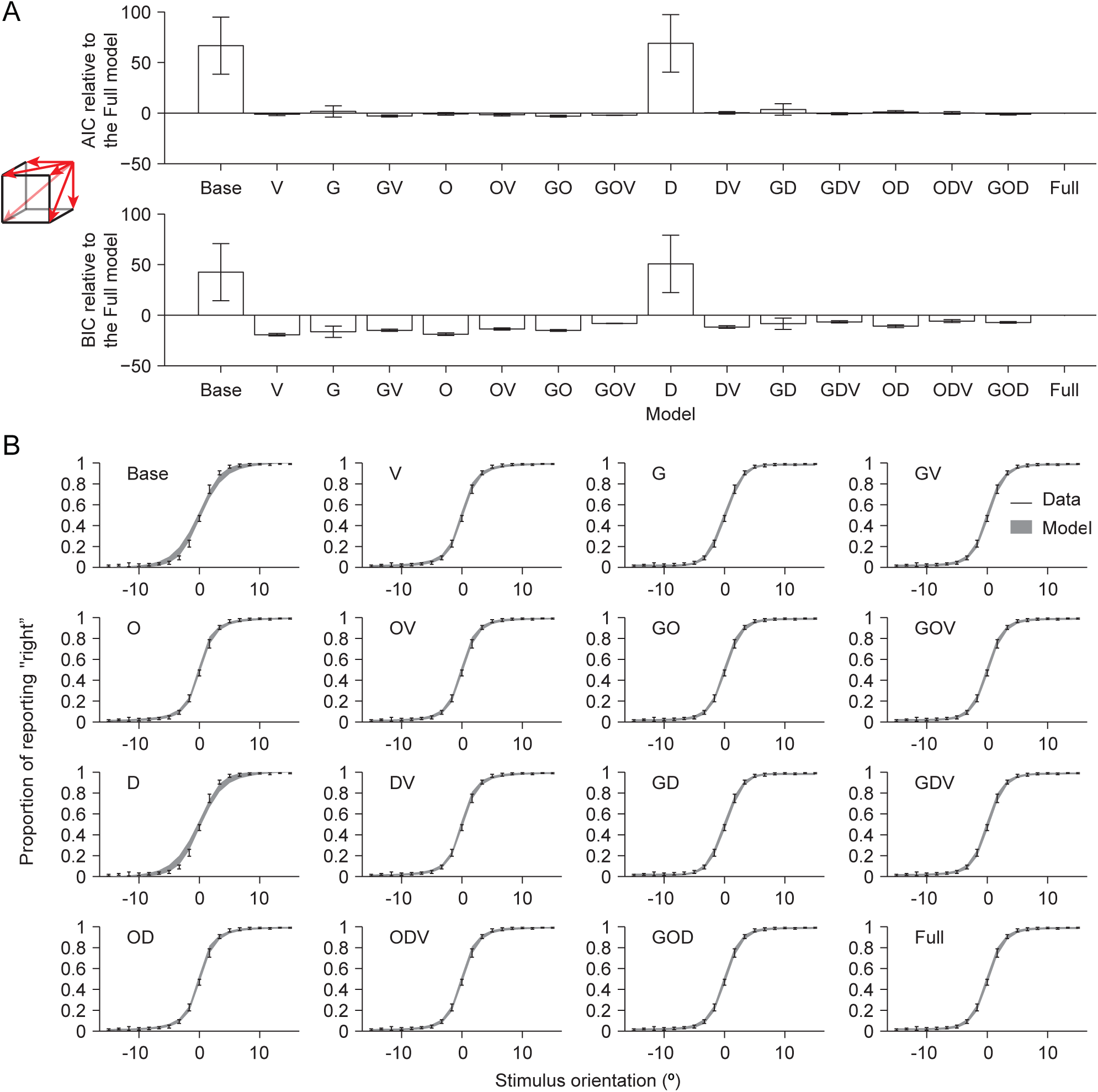
Experiment 3. **(A)** Complete model comparison. Mean and s.e.m. of the difference in AIC (top) and BIC (bottom) between each model and the Full model. **(B)** Proportion of reporting “right” as a function of target orientation: data and model fits.

**Figure A4.**
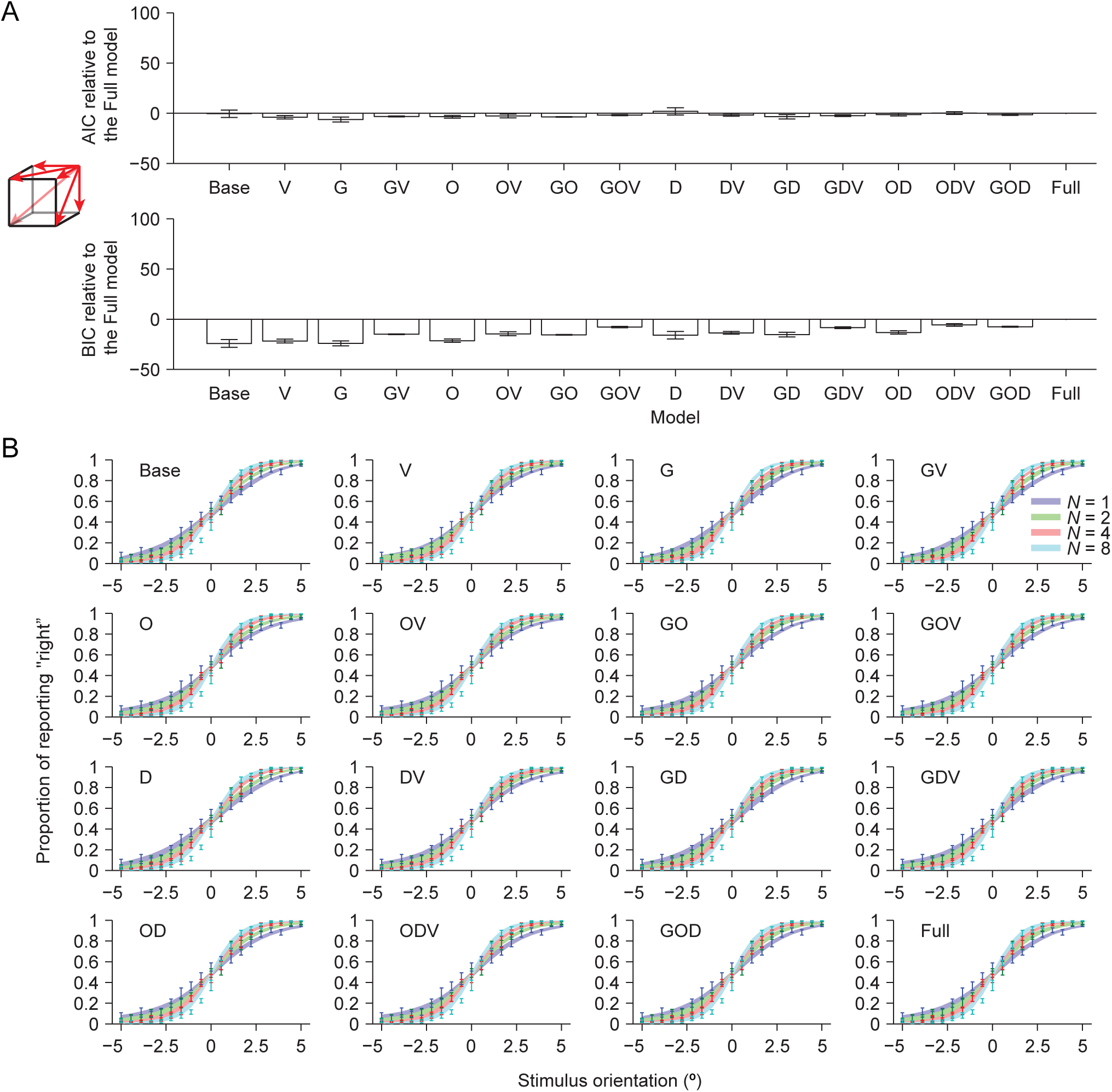
Experiment 4. **(A)** Complete model comparison. Mean and s.e.m. of the difference in AIC (top) and BIC (bottom) between each model and the Full model. **(B)** Proportion of reporting “right” as a function of set size and target orientation: data and model fits.

**Figure A5.**
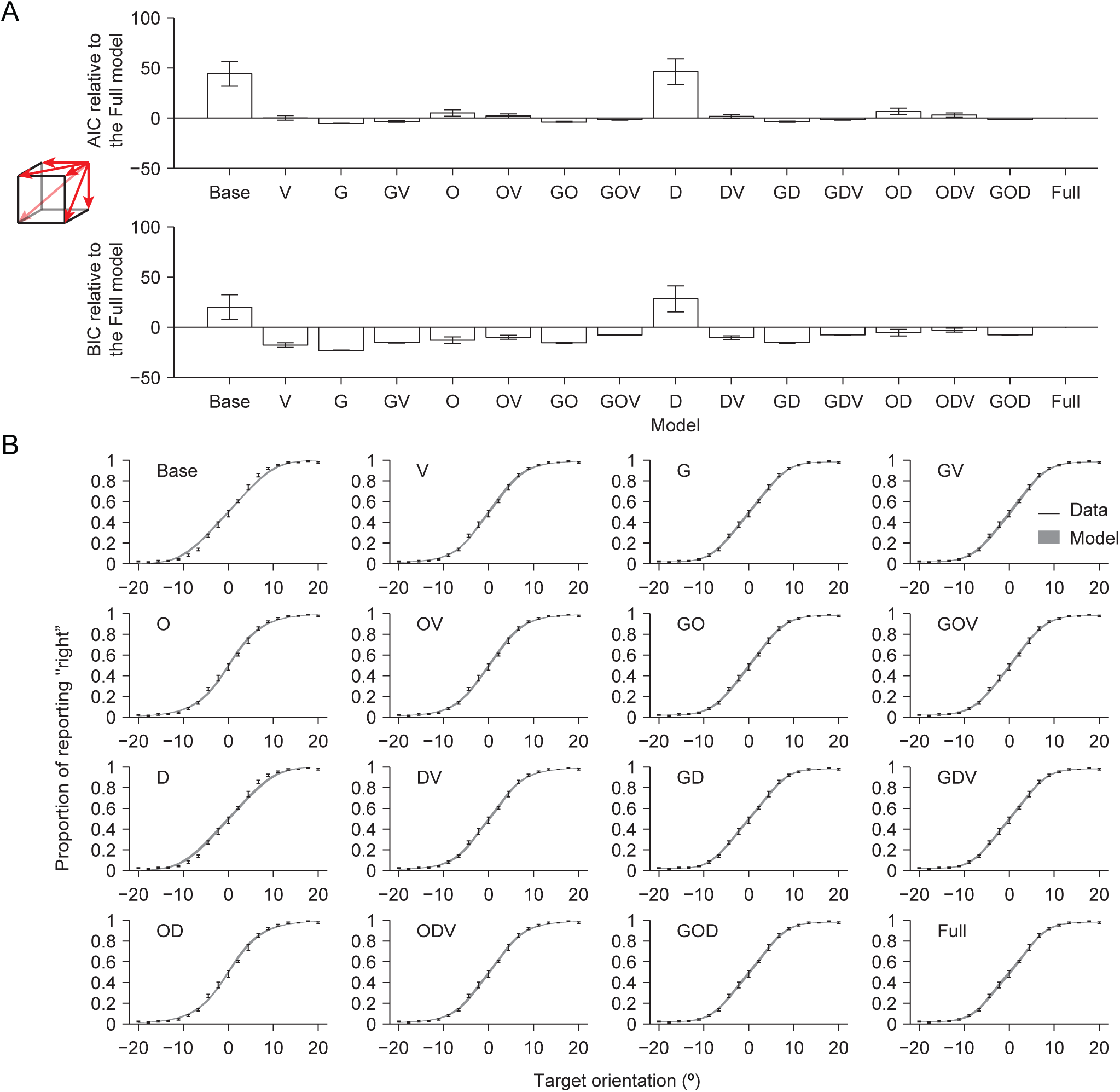
Experiment 5. **(A)** Complete model comparison. Mean and s.e.m. of the difference in AIC (top) and BIC (bottom) between each model and the Full model. **(B)** Proportion of reporting “right” as a function of target orientation.

**Figure A6.**
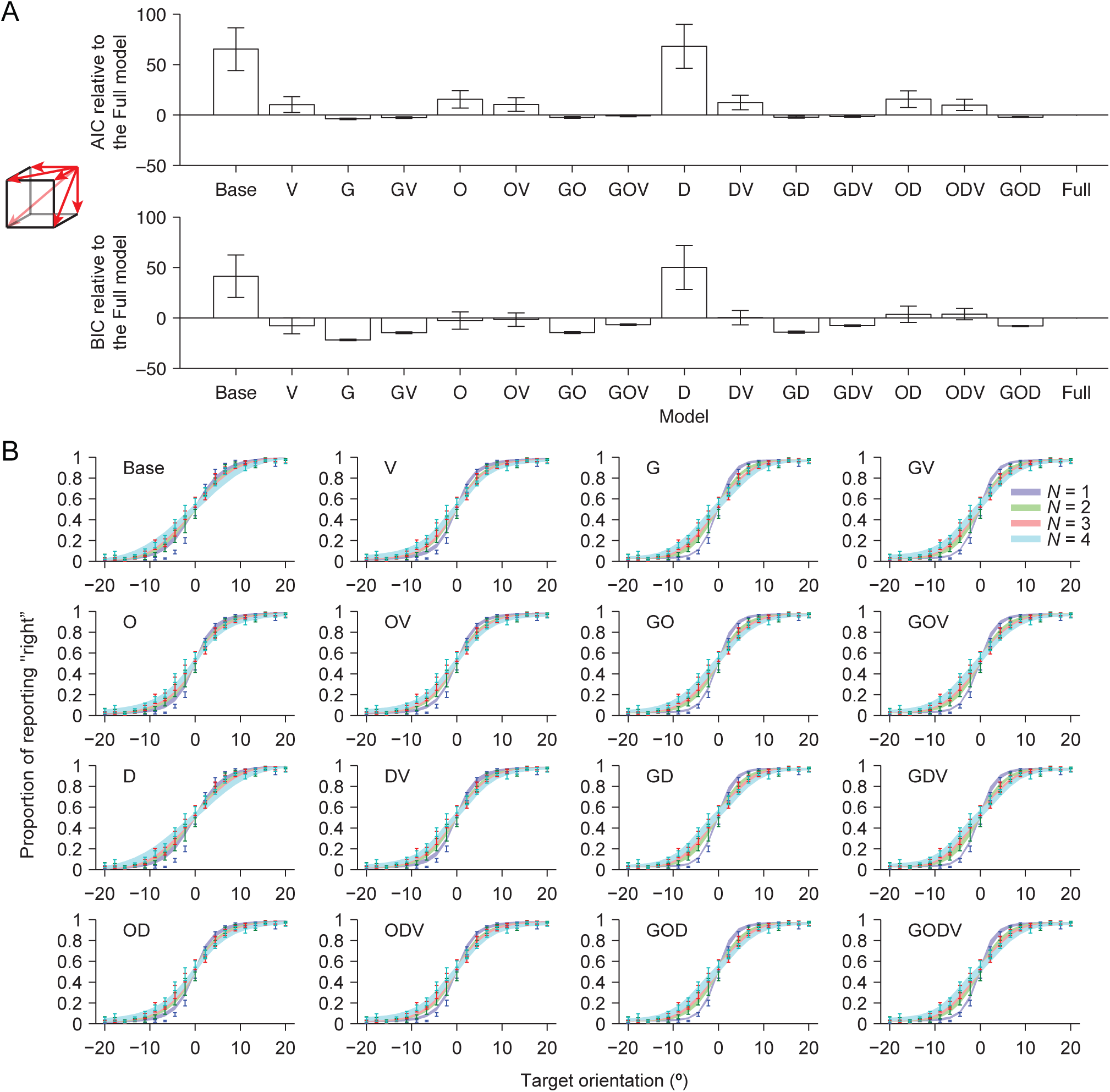
Experiment 6. **(A)** Complete model comparison. Mean and s.e.m. of the difference in AIC (top) and BIC (bottom) between each model and the Full model. **(B)** Proportion of reporting “right” as a function of set size and target orientation: data and model fits.

**Figure A7.**
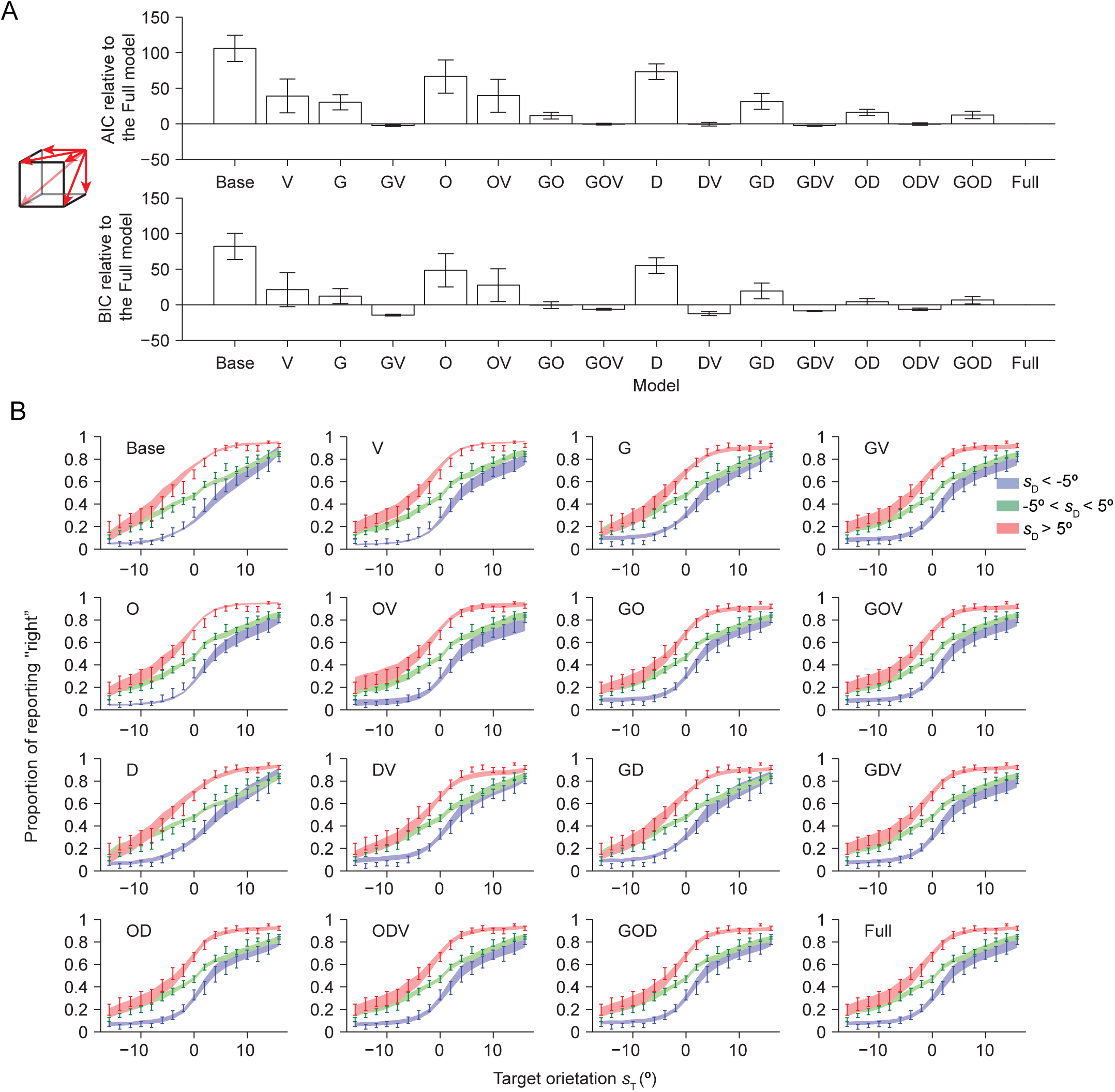
Experiment 7. **(A)** Complete model comparison. Mean and s.e.m. of the difference in AIC (top) and BIC (bottom) between each model and the Full model. **(B)** Proportion of reporting “right” as a function of target orientation: data and model fits.

**Figure A8.**
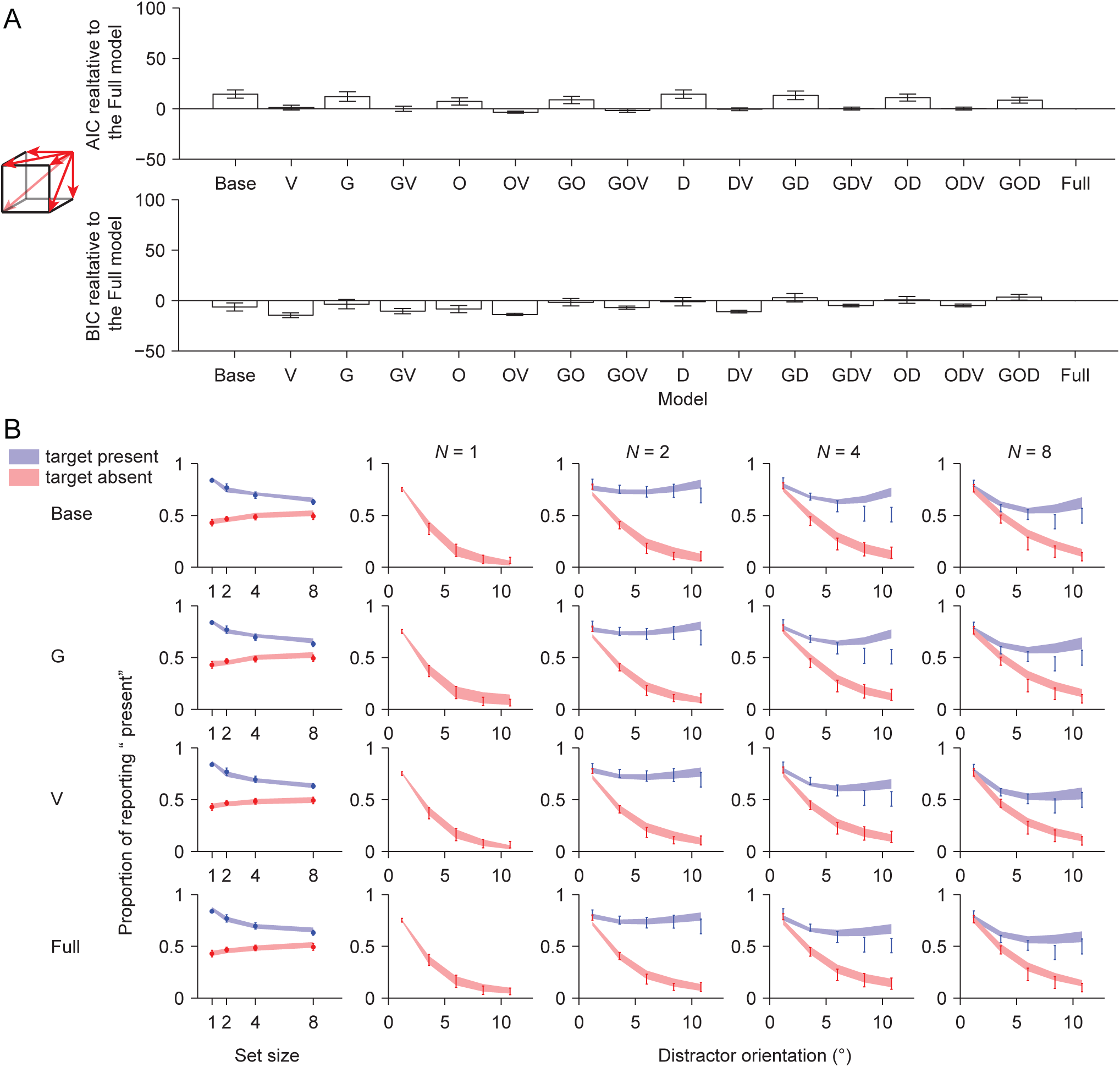
Experiment 8. **(A)** Complete model comparison. Mean and s.e.m. of the difference in AIC (top) and BIC (bottom) between each model and the Full model. **(B)** Proportion of reporting “present” as a function of set size, target presence, and the common orientation of the distractors: data and model fits.

**Figure A9.**
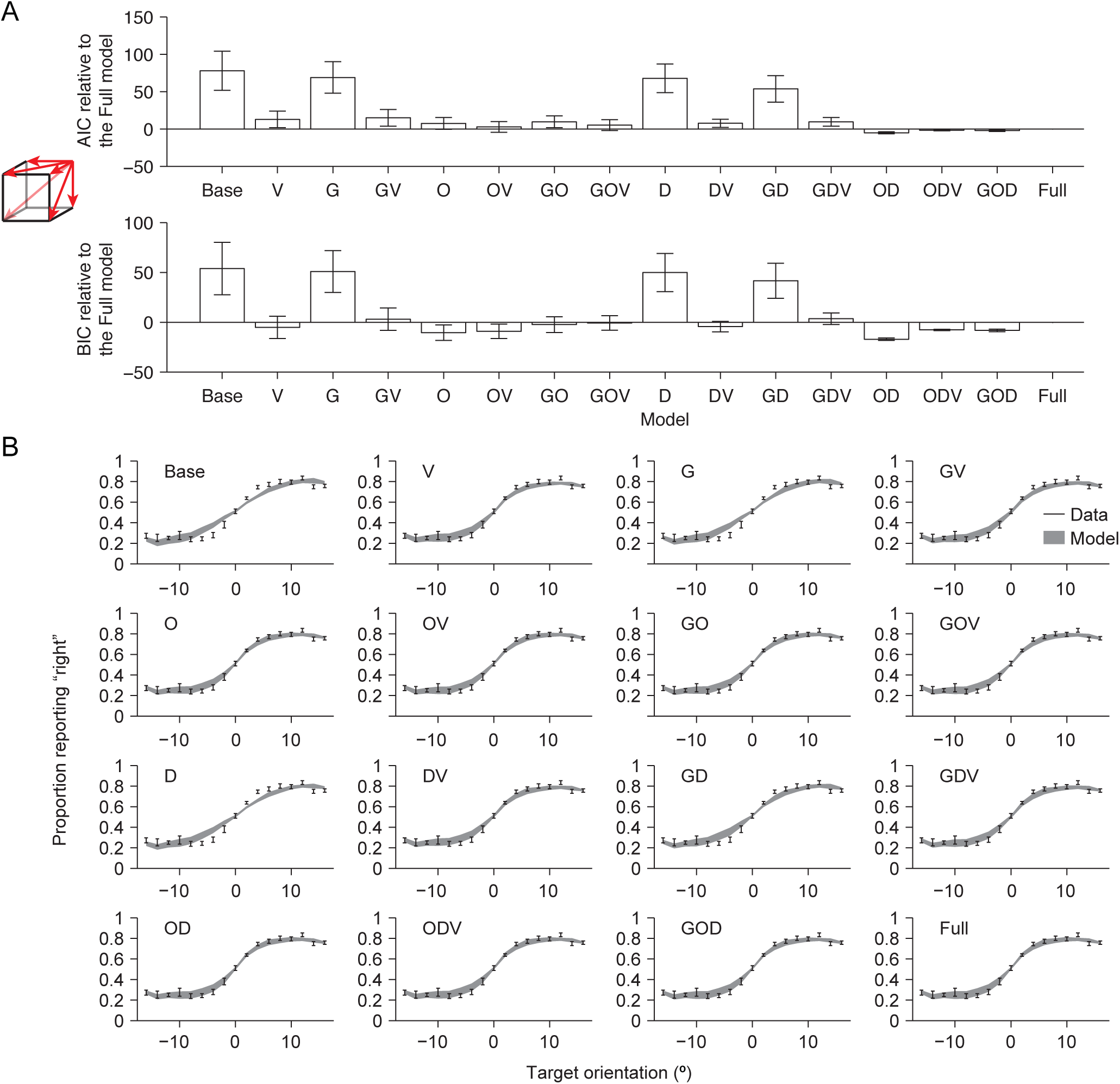
Experiment 9. **(A)** Complete model comparison. Mean and s.e.m. of the difference in AIC (top) and BIC (bottom) between each model and the Full model. **(B)** Proportion of reporting “right” as a function of target orientation.

**Figure A10.**
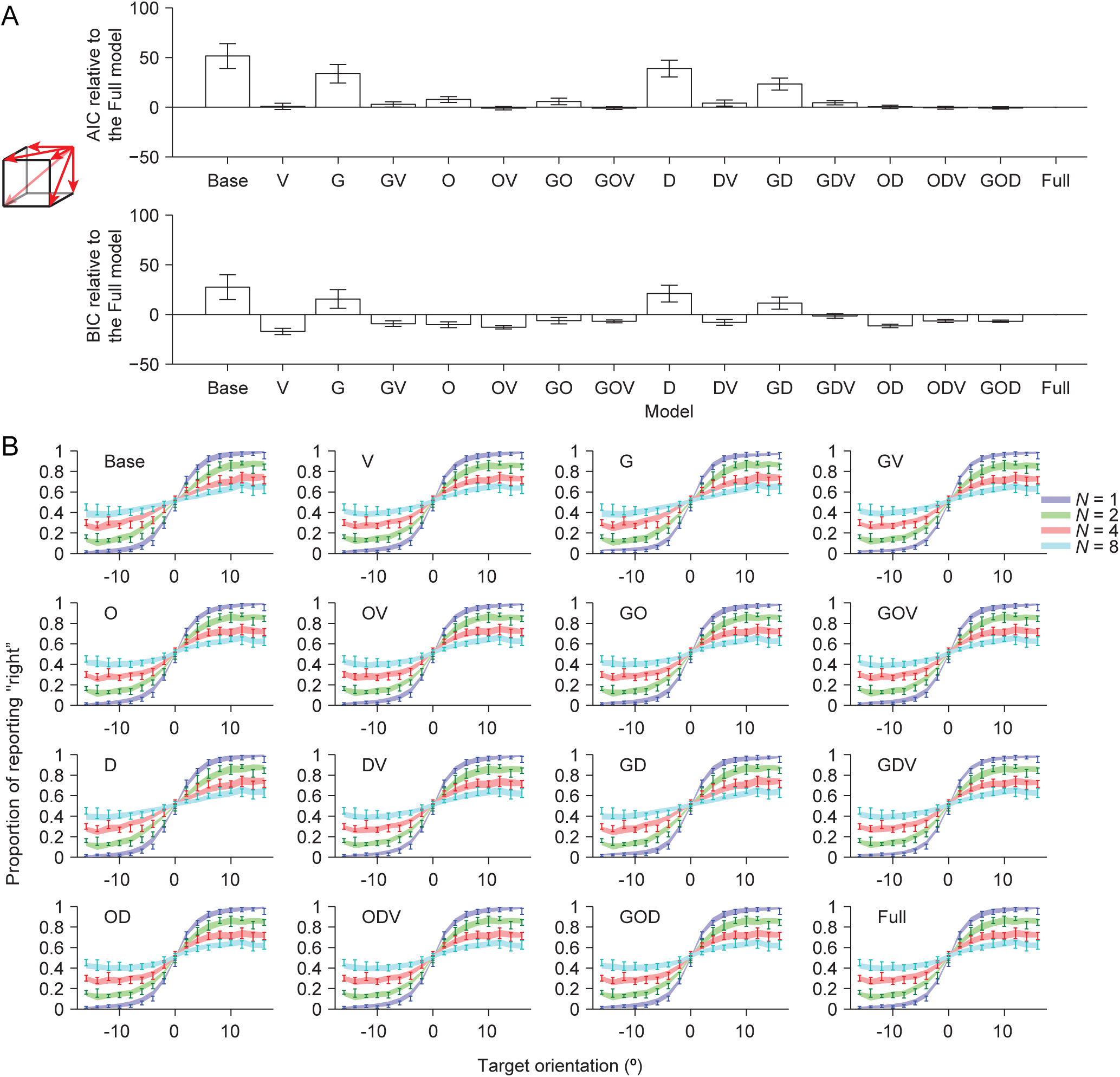
Experiment 10. **(A)** Complete model comparison. Mean and s.e.m. of the difference in AIC (top) and BIC (bottom) between each model and the Full model. **(B)** Proportion of reporting “right” as a function of target orientation.

**Figure A11.**
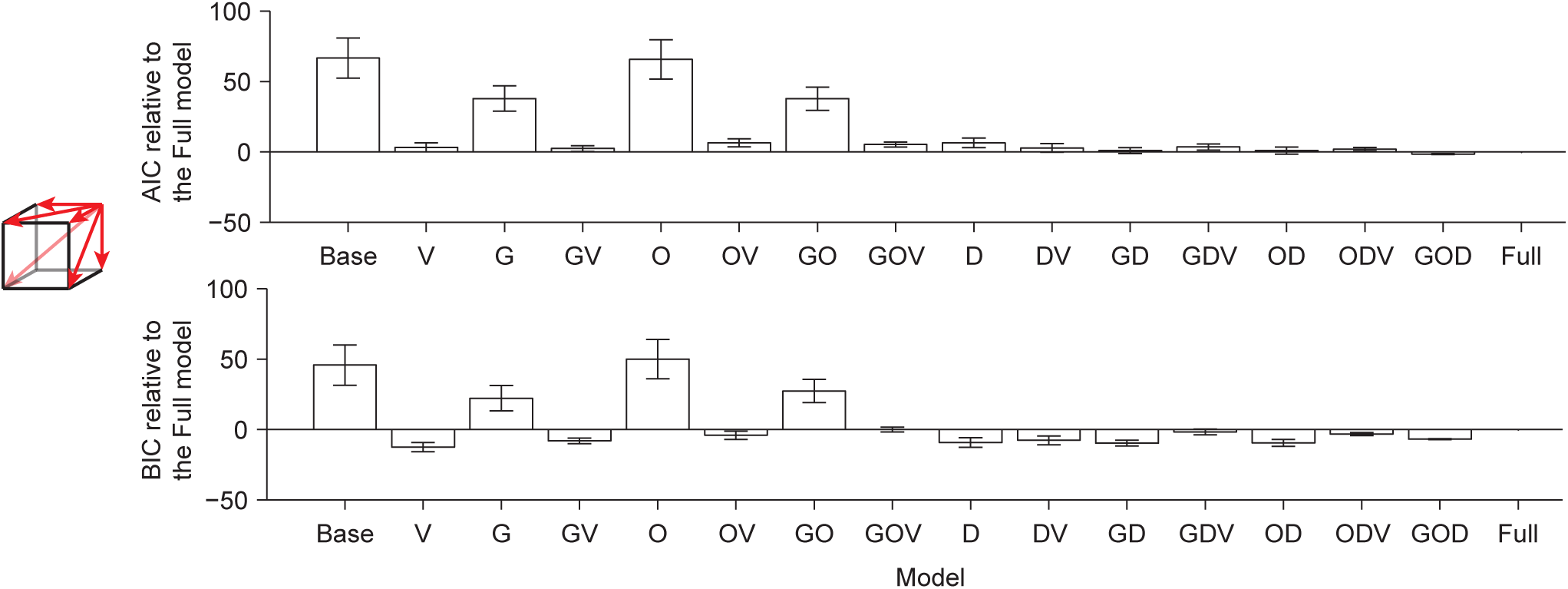
Complete model comparison in Experiment 11. Mean and s.e.m. of the difference in AIC (top) and BIC (bottom) between each model and the Full model.

In Experiments 7, 9, 10, and 11, mean KOD_AIC_(V) and mean KOD_BIC_(V) were both greater than 9.2, indicating very strong evidence that factor V is necessary to explain the data (**Figure 4B**). This is consistent with the model fits to the psychometric curves (**Figure 4E**, **Figures A7**, **A9** and **A10**, **panel B**, compare the G and GV models). In Experiment 8, KOD(V) was 12.1 ± 4.3 (AIC) and 6.9 ± 4.3 (BIC), indicating strong evidence that factor V is necessary (**Figure 4B**). In Experiments 1 to 6, however, mean KOD_AIC_(V) and mean KOD_BIC_(V) were both lower than 4.6, indicating little or no evidence that factor V is necessary. The big difference between KID(V) and KOD(V) in Experiments 1, 3, 5 and 6 arouse because factor G could also explain the data well, as indicated by a high KID(G) (**Figure 4A**) and illustrated by the model fits (**Figures 4D**, **A1**, **A3**, **A5**, **A6**).

Using LFLR, we found very strong evidence for the presence of factor V in Experiments 7, 9, 10, and 11, with mean 2·LFLP_AIC_(V) and mean 2·LFLP_BIC_(V) both greater than 9.2 (**Figure 4C**). We also found strong evidence for the presence of factor V in Experiment 8, with a 2·LFLP(V) equal to 12.7 ± 4.4 (AIC) and 7.8 ± 4.5 (BIC). The common feature of these five experiments was that distractors were variable across trials. In Experiments 7 and 8, the distractors were homogenous within a trial, while in Experiments 9 to 11, distractors were heterogeneous within a trial. We found little or no evidence for factor V in Experiments 1 to 6, with mean 2·LFLP_AIC_(V) and mean 2·LFLP_BIC_(V) both smaller than 4.6. The common feature of these experiments was that there were either no distractors (Experiments 1 to 4), or fixed distractors (Experiments 5 to 6).

Overall, with consideration of guessing, we found evidence for the presence of variable precision to be very little in Experiments 1 to 6, strong in Experiment 8, and very strong in Experiment 7, and 9 to 11.

How do these results compare to the visual short-term memory literature? The analog of a comparison between the G and V models has been made in several visual short-term memory experiments (D. T. Devkar et al., 2015; keshvari et al., 2012, 2013; van den Berg et al., 2012), with V fitting better. In one paper, GV was compared to G, with GV fitting better (Fougnie et al., 2012; van den Berg et al., 2014). Finally, in one paper, a form of GV (there called VP-F) was compared to both V (VP-A) and a form of G (EP-F), with GV fitting best; however, the guessing was set size-dependent in a specific way (dictated by an item limit) (Fougnie et al., 2012; van den Berg et al., 2014). All these results were obtained in experiments with multiple set sizes and “heterogeneous distractors”, most similar to our Experiments 10 and 11. Indeed, the results are consistent with ours in those experiments.

### The importance of variable precision (V) when taking into account guessing (G) and the oblique effect (O)

Although we found evidence for variable precision (V) in Experiments 7 to 11 when considering G, some of the variability could be explained by other confounding factors. We first considered the oblique effect (O), the phenomenon that oblique orientations are encoded with lower precision than cardinal ones (Appelle, 1972; Girshick et al., 2011; Pratte et al., 2017). We implemented O using a rectified sine function (**Figure 2B** and **Eq. (2)**).

Inclusion of factor O did not qualitatively change the importance of factor V in Experiments 1 to 8 and 11 (**Figure 5A-C**). However, it greatly reduced the importance of factor V in Experiments 9 and 10. KOD(V) was 4.4 ± 3.3 (AIC) and −1.6 ± 3.3 (BIC) in Experiment 9, and 6.7 ± 2.7 (AIC) and 0.7 ± 2.7 (BIC) in Experiment 10 (**Figure 5B**), indicating little or no evidence that factor V is necessary. Consistently, 2·LFLP(V) was 5.4 ± 3.5 (AIC) and 0.9 ± 4.4 (BIC) in Experiment 9 (**Figure 5C**), indicating little or no evidence for the presence of factor V. 2·LFLP(V) was 8.5 ± 2.9 (AIC) and 6.4 ± 3.3 (BIC) in Experiment 10 (**Figure 5C**), indicating moderate evidence for the presence of factor V.

**Figure 5.**
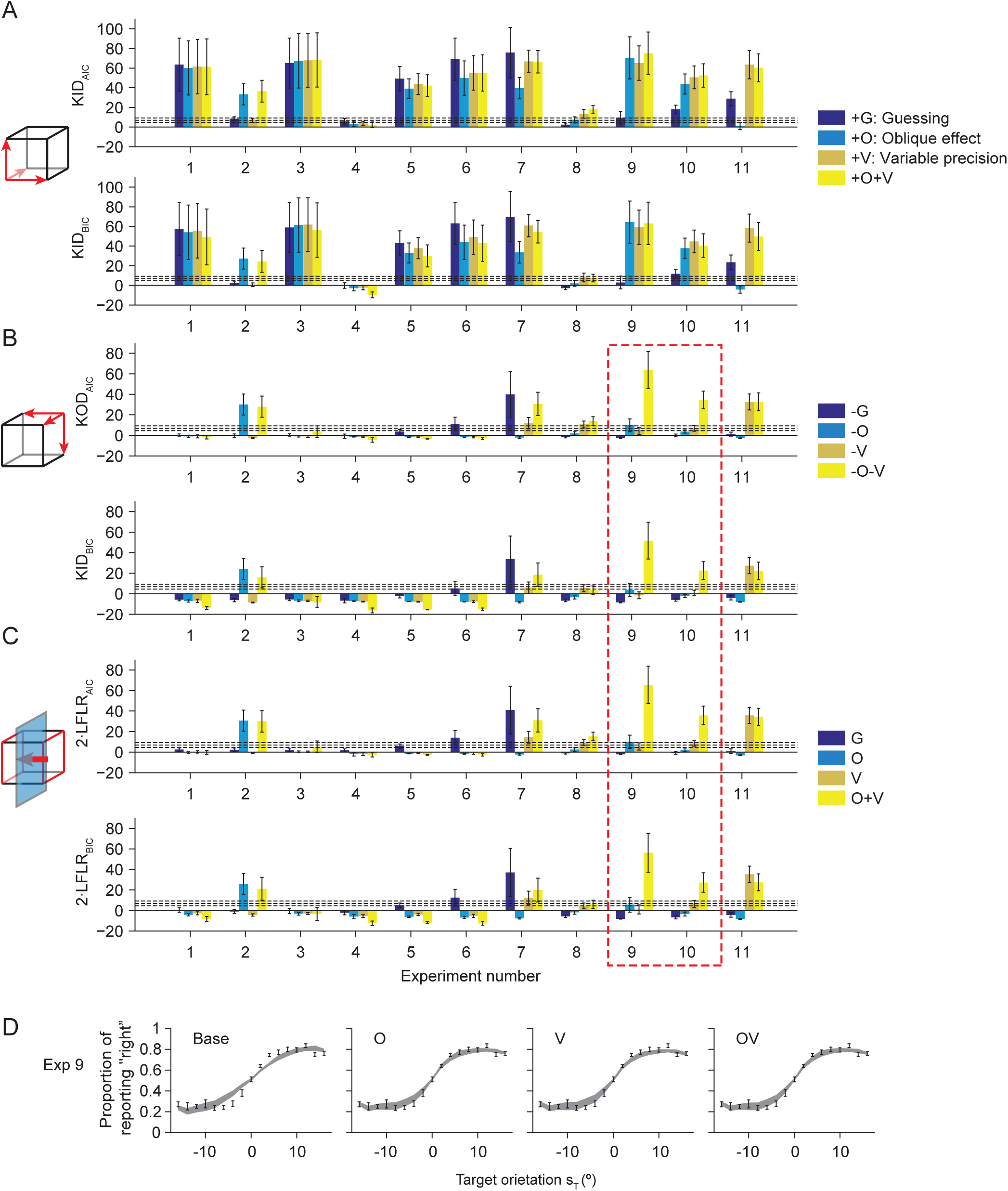
Factor importance among guessing (G), oblique effect (O), and residual variable precision (V). The red dashed box marks the major changes (compared to Fig. 4) in the evidence for the importance of factor V when taking factor O into consideration. **(A-C)** Mean and s.e.m. of KID (A), KOD (B), and 2-LFLR (C) based on AIC (top) or BIC (bottom) for the factors G, O, V, and the OV combination, in all experiments. **(D)** Model fits in Experiment 9. The O, V, and OV models fit the data almost equally well, and better than the Base model.

In Experiments 9 and 10, evidence for the presence of factor V was reduced because factor O could also explain the data well. KID(O) was 70 ± 21 (AIC) and 64 ± 21 (BIC) in Experiment 9, and 44 ± 10 (AIC) and 38 ± 10 (BIC) in Experiment 10 (**Figure 5A**). This can also be seen in the model fits of model O (**Figure 5D**). However, knocking out factor O did not cause large changes in AIC or BIC: KOD(O) was 9.8 ± 6.3 (AIC) and 3.8 ± 6.3 (BIC) in Experiment 9, and 3.7 ± 1.5 (AIC) and −2.3 ± 1.5 (BIC) in Experiment 10 (**Figure 5B**), indicating little or no evidence that factor O is necessary. knocking out both factors O and V was disastrous, resulting in a KOD(OV) of 64 ± 18 (AIC) and 52 ± 18 (BIC) in Experiment 9, and of 34.5 ± 8.7 (AIC) and 22.5 ± 8.7 (BIC) in Experiment 10 (**Figure 5B**). This “non-linear” phenomenon seemed to occur because factors O and V could stand in for each other in explaining the data, therefore neither was necessary, but having at least one of them was important. This is consistent with the model fits to the psychometric curves (**Figure 5D**, **Figures A9B**, **A10B**). Consistently, 2·LFLR was not high for either factor O or factor V, but was high for their combination: 65 ± 18 (AIC) and 56 ± 19 (BIC) in Experiment 9, and 36.9 ± 9.0 (AIC) and 27.2 ± 9.4 (BIC) in Experiment 10 (**Figure 5C**).

In the visual short-term memory literature, Pratte et al. compared models similar to G, V, GV, GO, OV, and GOV (Pratte et al., 2017); again however, the guessing was set size-dependent in a specific way (dictated by an item limit). They found that V fitted better than G but worse than GV. Adding factor O flipped the first result: GO fitted better than OV; however, both still fitted much worse than GOV. Although they did not test the Base and O models, this last result suggests evidence for the presence of V. The experiments by Pratte et al. were again most similar to our Experiments 10 and 11, featuring multiple set sizes and heterogeneous distractors. In Experiment 10, we also found that accounting for factor O changed our evidence for the presence of factor V (V fitted better than G but GO fitted as well as OV), and (using LFLR) we found moderate evidence for the presence of V. In Experiment 11, considering factor O did not change our evidence for the importance of factor V, and we still found strong evidence for the presence of factor V.

### The importance of variable precision (V) when taking into account guessing (G), the oblique effect (O), and decision noise (D)

Besides guessing and the oblique effect, another confounding factor might be noise in the decision stage (or suboptimal inference, which can look like decision noise). We examined evidence for the importance of V when accounting for guessing (G), the oblique effect (O), and decision noise (D). We modeled D as Gaussian noise added to the log posterior ratio (**Eq. (3)**).

In Experiments 1 to 10, the inclusion of factor D did not change the evidence for the importance of factor V much (**Figure 6A-C**). In Experiment 8, evidence for the presence of factor V changed from strong to moderate (**Figure 6B**). In Experiment 10, the inclusion of factor D slightly reduced the evidence for the presence of factor V. 2·LFLP(V) was 2.3 ± 2.3 (AIC) and 1.9 ± 3.3, indicating little or no evidence for the presence of factor V.

**Figure 6.**
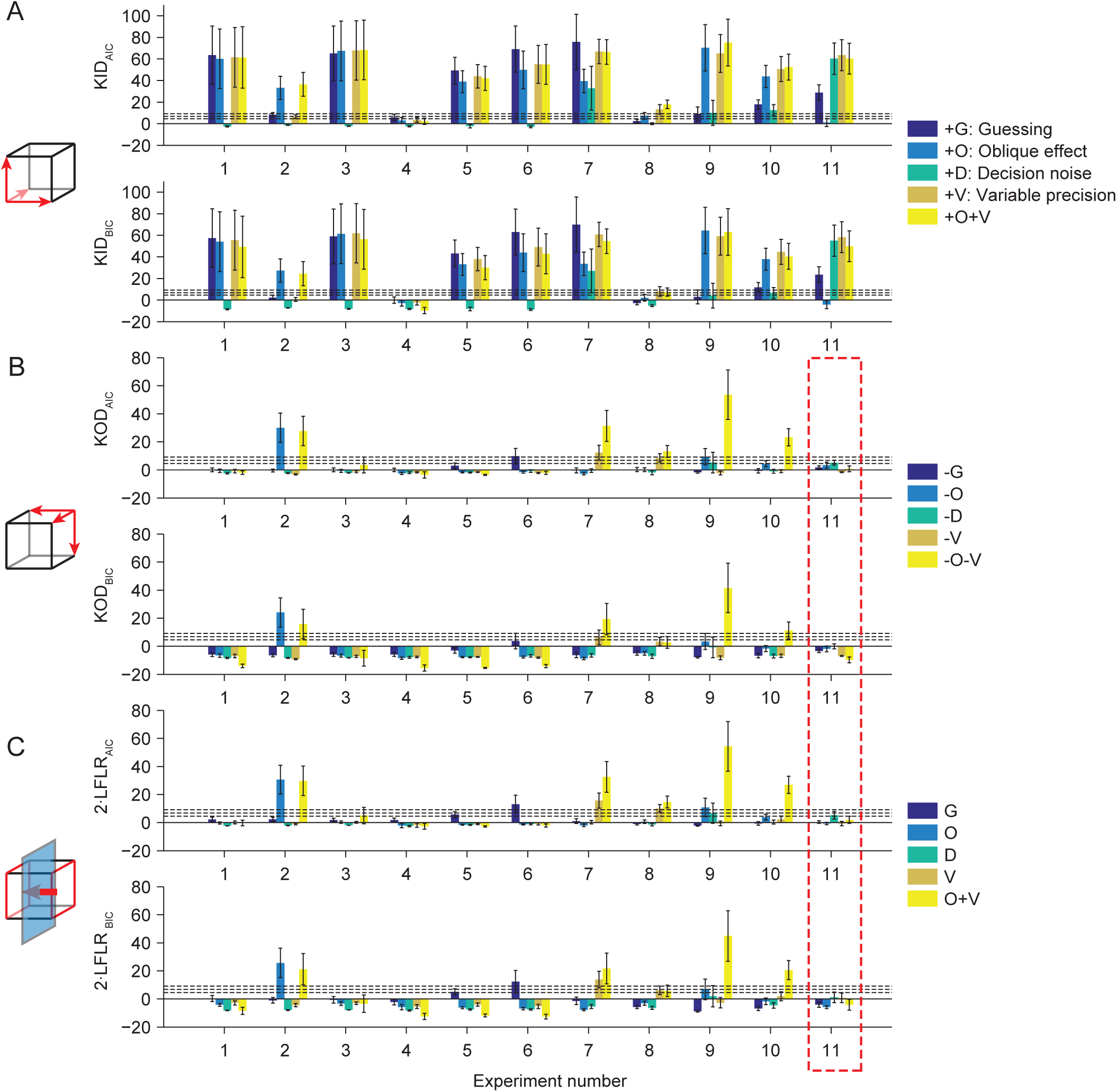
Factor importance: guessing (G), oblique effect (O), decision noise (G) and the residual variable precision (V). The red dashed box marks the major changes (compared to Fig. 5) in the evidence for the importance of factor V when taking factor D into consideration. **(A-C)** Mean and s.e.m. of KID (A), KOD (B), and 2-LFLR (C) based on AIC (top) or BIC (bottom) for the factors G, O, D, V, and the OV combination, in all experiments.

In Experiment 11, however, the inclusion of D greatly reduced the importance of factor V. KOD_AIC_(V) and KOD_BIC_(V) were negative (**Figure 6B**), indicating no evidence that factor V is necessary. Consistently, 2·LFLP(V) was −0.6 ± 1.6 (AIC) and 0.7 ± 3.4 (BIC) in Experiment 11 (**Figure 6C**), indicating little or no evidence for the presence of factor V. The reason is probably that factor D can also explain the data: KID(D) was 60 ± 14 (AIC) and 55 ± 14 (BIC) (**Figure 6A**), consistent with the model fits of the D model (**Figure 7**). However, knocking out factor D did not cause large KODs (**Figure 6B**), suggesting that factor D is useful, but not necessary.

**Figure 7.**
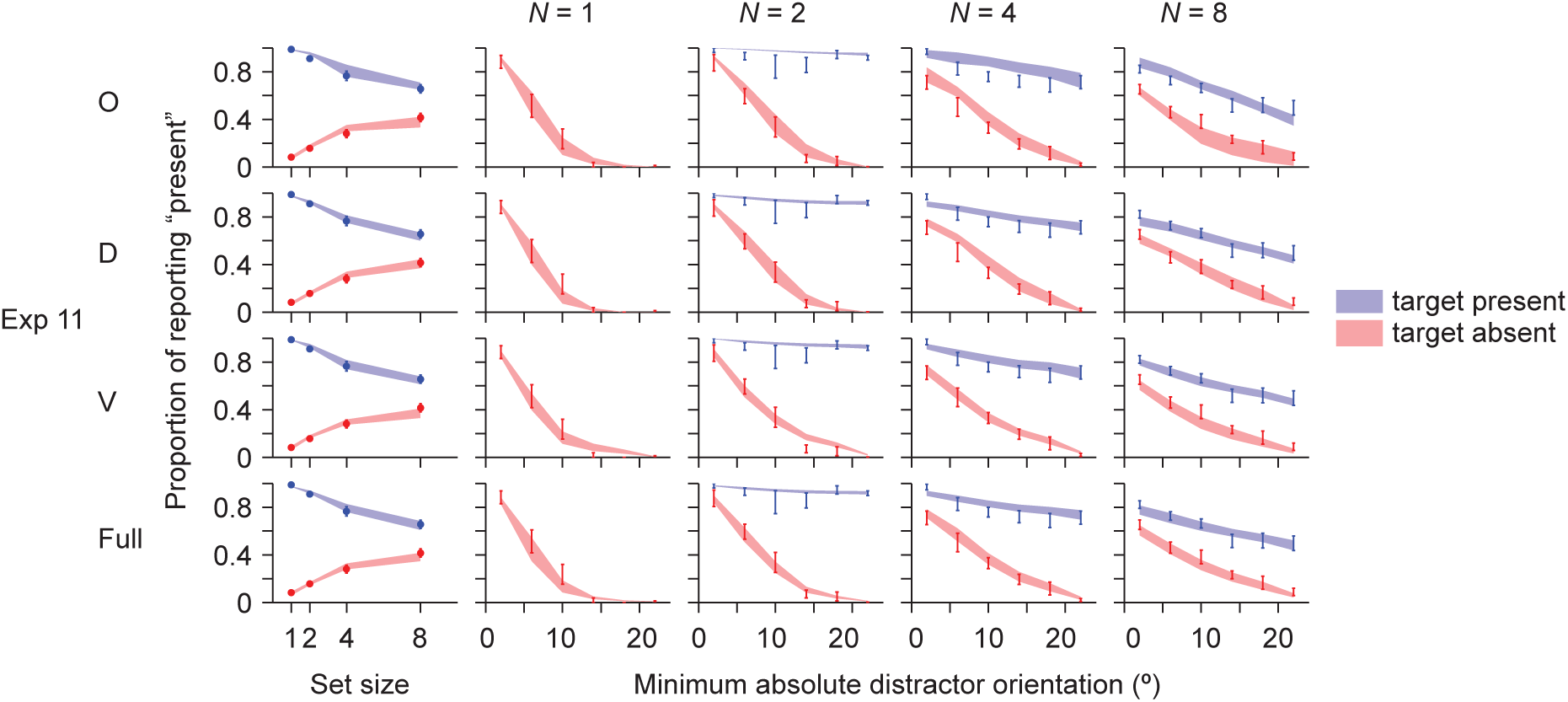
Model fits in Experiment 11. Proportion of reporting “target present” as a function of set size, target presence, and the smallest circular distance in orientation space between the target and any of the distractors. The D and V models fit the data almost as well as the Full model, and better than the O model.

In summary, when accounting for guessing, the oblique effect, and decision noise, we only found very strong evidence for the presence of the residual variable precision in Experiment 7. Experiment 7 was the only orientation categorization task in which the distractors were homogeneous but varied across trials. In Experiment 8, which was a target detection task also with homogeneous variable distractors, we found moderate evidence for residual variable precision, suggesting that homogeneous variable distractors might induce residual variable precision.

To our knowledge, no previous studies have compared models containing all four factors G, O, D, and V.

### Relationship between task features and importance of factors other than the residual variable precision

The variation of our experiments also enables us to relate the features of tasks to the importance of factors other than the residual variable precision. The experiments differed in the following design features (**Table 4**): set size greater than 1 (divided attention), set size variability, number of targets greater than 1, task type (categorization or detection), the distribution of the target orientation, the distribution of the orientation of the reference (Experiment 2) or the distractors (all other experiments), distractor variability across displays, distractor variability within displays, and the presence of ambiguity (in the form of overlapping target-distractor category distributions).

**Table 4:**
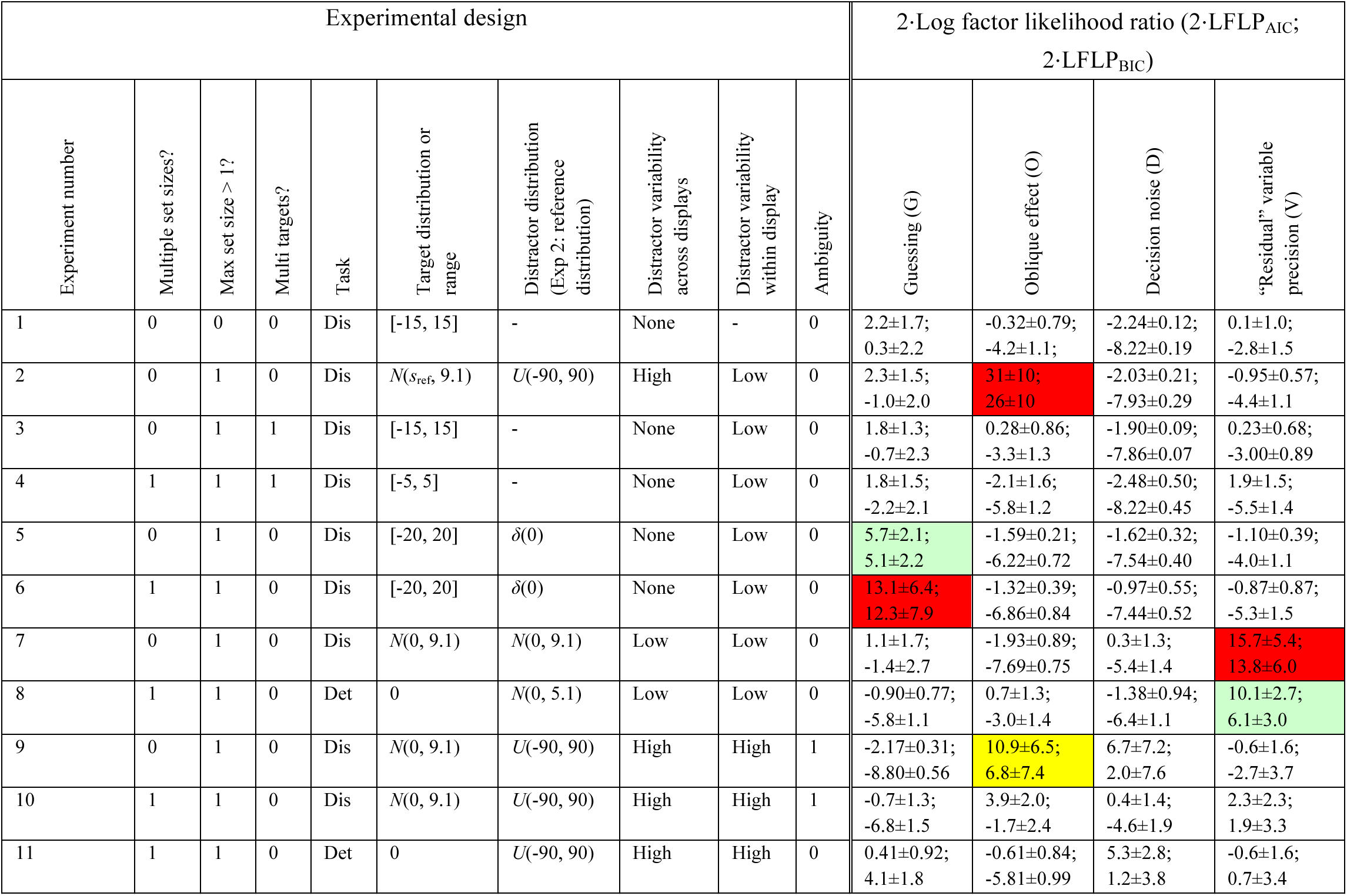
Features of experiments and the evidence for the presence of factors. Here are the abbreviations and notations. Dis: Discrimination; Det: Detection; ref: reference; *s*_ref_: reference orientation; *U*(*a, b*) denotes a continuous uniform distribution on the interval [*a, b*]; *N*(*s*_0_, *σ*) denotes a Gaussian distribution with a mean of *s*_0_ and a standard deviation of *σ*; *δ*(*a*) denotes a Dirac delta function at *a*. The unit of orientation is degrees, and we “converted” the (highly concentrated) Von Mises distributions to Gaussian distributions to make the comparison across experiments easier. Red, yellow and light green mark the very strong, strong, and moderate evidence for the presence of a factor, respectively, based on our criteria in **Table 3b**.

| Experimental design |  |  |  |  |  |  |  |  |  | 2·Log factor likelihood ratio (2·LFLP<sub>AIC</sub>;<br>2·LFLP<sub>BIC</sub>) |  |  |  |
| --- | --- | --- | --- | --- | --- | --- | --- | --- | --- | --- | --- | --- | --- |
| Experiment number | Multiple set sizes? | Max set size > 1? | Multi targets? | Task | Target distribution or range | Distractor distribution (Exp 2: reference distribution) | Distractor variability across displays | Distractor variability within display | Ambiguity | Guessing (G) | Oblique effect (O) | Decision noise (D) | “Residual” variable precision (V) |
| 1 | 0 | 0 | 0 | Dis | [-15, 15] | - | None | - | 0 | 2.2±1.7;<br>0.3±2.2 | -0.32±0.79;<br>-4.2±1.1; | -2.24±0.12;<br>-8.22±0.19 | 0.1±1.0;<br>-2.8±1.5 |
| 2 | 0 | 1 | 0 | Dis | $N(s_{\text{ref}}, 9.1)$ | $U(-90, 90)$ | High | Low | 0 | 2.3±1.5;<br>-1.0±2.0 | 31±10;<br>26±10 | -2.03±0.21;<br>-7.93±0.29 | -0.95±0.57;<br>-4.4±1.1 |
| 3 | 0 | 1 | 1 | Dis | [-15, 15] | - | None | Low | 0 | 1.8±1.3;<br>-0.7±2.3 | 0.28±0.86;<br>-3.3±1.3 | -1.90±0.09;<br>-7.86±0.07 | 0.23±0.68;<br>-3.00±0.89 |
| 4 | 1 | 1 | 1 | Dis | [-5, 5] | - | None | Low | 0 | 1.8±1.5;<br>-2.2±2.1 | -2.1±1.6;<br>-5.8±1.2 | -2.48±0.50;<br>-8.22±0.45 | 1.9±1.5;<br>-5.5±1.4 |
| 5 | 0 | 1 | 0 | Dis | [-20, 20] | $\delta(0)$ | None | Low | 0 | 5.7±2.1;<br>5.1±2.2 | -1.59±0.21;<br>-6.22±0.72 | -1.62±0.32;<br>-7.54±0.40 | -1.10±0.39;<br>-4.0±1.1 |
| 6 | 1 | 1 | 0 | Dis | [-20, 20] | $\delta(0)$ | None | Low | 0 | 13.1±6.4;<br>12.3±7.9 | -1.32±0.39;<br>-6.86±0.84 | -0.97±0.55;<br>-7.44±0.52 | -0.87±0.87;<br>-5.3±1.5 |
| 7 | 0 | 1 | 0 | Dis | $N(0, 9.1)$ | $N(0, 9.1)$ | Low | Low | 0 | 1.1±1.7;<br>-1.4±2.7 | -1.93±0.89;<br>-7.69±0.75 | 0.3±1.3;<br>-5.4±1.4 | 15.7±5.4;<br>13.8±6.0 |
| 8 | 1 | 1 | 0 | Det | 0 | $N(0, 5.1)$ | Low | Low | 0 | -0.90±0.77;<br>-5.8±1.1 | 0.7±1.3;<br>-3.0±1.4 | -1.38±0.94;<br>-6.4±1.1 | 10.1±2.7;<br>6.1±3.0 |
| 9 | 0 | 1 | 0 | Dis | $N(0, 9.1)$ | $U(-90, 90)$ | High | High | 1 | -2.17±0.31;<br>-8.80±0.56 | 10.9±6.5;<br>6.8±7.4 | 6.7±7.2;<br>2.0±7.6 | -0.6±1.6;<br>-2.7±3.7 |
| 10 | 1 | 1 | 0 | Dis | $N(0, 9.1)$ | $U(-90, 90)$ | High | High | 1 | -0.7±1.3;<br>-6.8±1.5 | 3.9±2.0;<br>-1.7±2.4 | 0.4±1.4;<br>-4.6±1.9 | 2.3±2.3;<br>1.9±3.3 |
| 11 | 1 | 1 | 0 | Det | 0 | $U(-90, 90)$ | High | High | 0 | 0.41±0.92;<br>4.1±1.8 | -0.61±0.84;<br>-5.81±0.99 | 5.3±2.8;<br>1.2±3.8 | -0.6±1.6;<br>0.7±3.4 |

By examining the importance of factors in all these 11 experiments, we found that some factors were important when certain features were present. We now summarize the evidence for the importance of each factor across experiments and attempt to make a connection to the features of the experiments; **Table 4** lists the evidence for the presence of each factor (2·LFLR) in each experiment.

#### Guessing (G)

Consistent with the notion that guessing is widespread in psychophysical tasks, we found that in many experiments (Experiments 1, 3, 5, 6, 7), KID(G) was greater than 9.2 (**Figure 6A**) and the G model provided clearly better fits to the psychometric curves than the Base model (**Figure 8A**, **Figures A1**, **A3**, **A5**, **A6**, **A7**, **panel B**). Among these experiments, in Experiment 6, factor G was necessary (**Figure 6B**) and had 2·LFLR of 13.1 ± 6.4 (AIC) and 12.3 ± 7.9 (BIC) (**Figure 6C**), indicating very strong evidence for the presence of factor G. In this experiment, the target orientation took values between −20° and 20°, which was the largest range among all experiments. Moreover, set size could be low (1 or 2). A mistake on a trial with low set size and a strongly tilted target could only be explained by guessing. In Experiment 5, in which the stimulus range was the same but set size was equal to 4, factor G was no longer necessary, but 2·LFLR(G) was 5.7 ± 2.1 (AIC), and 5.1 ± 2.2 (BIC), respectively, indicating moderate evidence for the presence of factor G. Across all experiments, it seems that the larger the proportion of easy trials, the higher the evidence for the importance of factor G. With fewer easy trials, models without factor G fitted the data equally well as models with factor G, by estimating a lower encoding precision (**Figure 8A**). For example, in Experiment 4, where the target orientation range was narrow (between −5° and 5°), the Base model fitted as well as the G model, but the estimated precision was lower.

**Figure 8.**
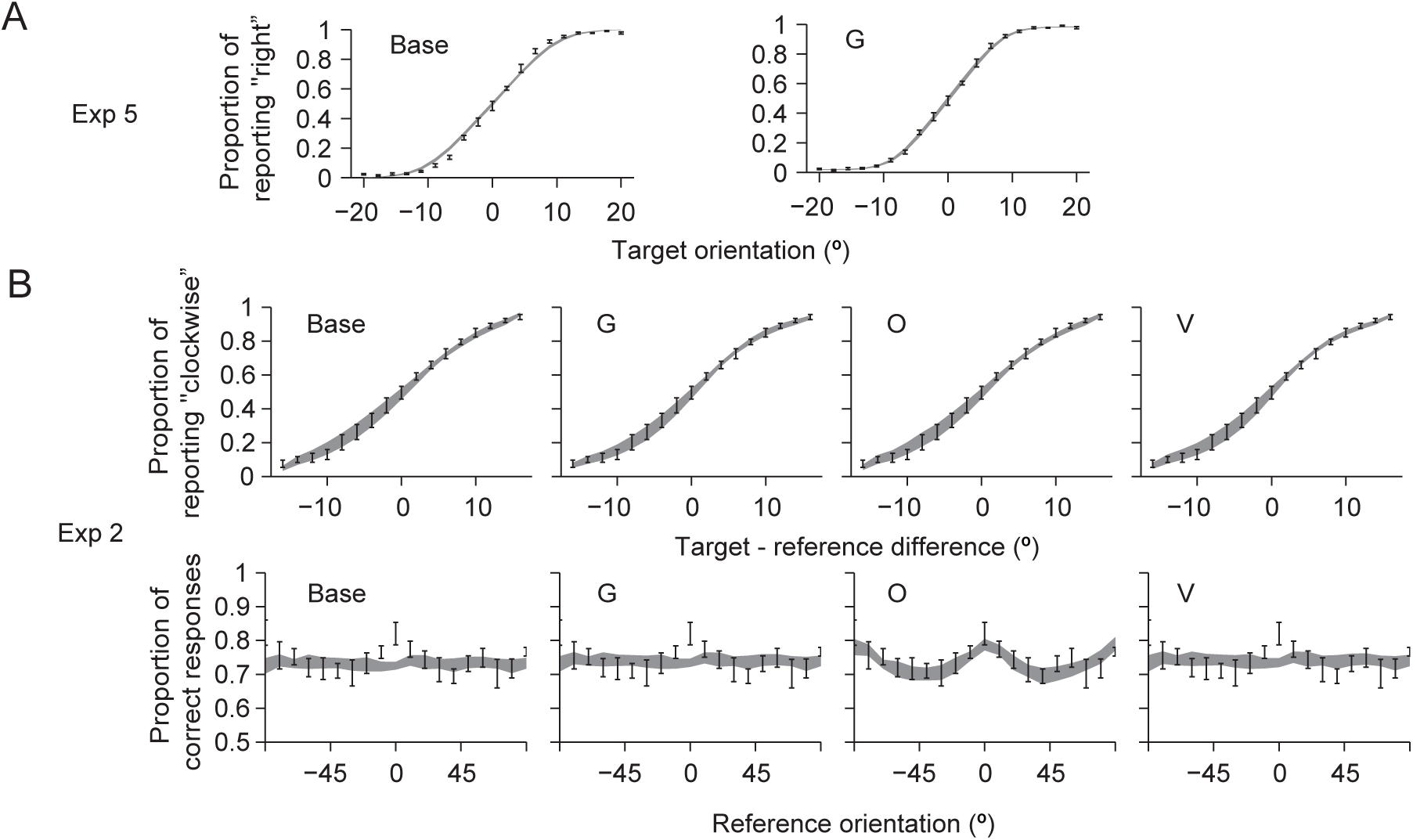
Model fits show that factor G and factor O are important in some experiments. **(A)** Model fits in Experiment 5 show the importance for factor G. The G model fits better than the Base model. **(B)** Model fits in Experiment 2 show the importance of factor O. Top: Proportion of reporting “clockwise” as a function of the orientation difference between the target and the reference, collapsed across reference orientations. Bottom: Proportion of reporting “clockwise” as a function of the reference orientation, collapsed across target orientations. The O model fits better than the Base, G, and V models.

#### Oblique effect (O)

We expected that O would be easier to detect when the stimulus distribution covered a larger orientation range. Indeed, we found very strong evidence for the presence of factor O in Experiment 2 and strong evidence in Experiment 9 (**Figure 6C**), in which the stimulus distribution covered the entire orientation space (**Figure 1**, **Experiment 2, Experiment 9**). This can also be seen in the model fits (**Figure 8B**). In Experiments 10 and 11, however, although the distractor distribution also covered the entire space, the evidence for factor O was weak. In Experiment 10, this might be because the experiment also contained a set size 1 condition, in which there were no distractors. In Experiment 11, stimuli with large tilts were informative of factor O, but because of the task structure, these stimuli were weighted less in the optimal decision rule (**Appendix 1, Experiment 11**), therefore perhaps making factor O harder to detect. Furthermore, because strongly tilted stimuli were less relevant to the task, subjects may have paid less attention to them. The weak evidence for the presence of factor O is consistent with previous findings that the oblique effect is weaker when stimuli are unattended (kelly & Matthews, 2011; Takács, Sulykos, Czigler, Barkaszi, & Balázs, 2013).

#### Decision noise (D)

Decision noise might reflect random variability or systematic suboptimality in the decision stage (Beck et al., 2012). We found little or no evidence for the presence of factor D in any of our experiments (**Figure 6C**). This suggests that human subjects are close to optimal. This is consistent with the conclusion of our previous paper (Shen & Ma, 2016), where we compared many suboptimal decision rules with the optimal rule in an orientation categorization task (Experiment 7 in this paper) and found that the more similar a suboptimal rule was to the optimal rule, the better it fitted the data.

### Causes for false negatives

Overall, we were conservative when claiming evidence for a certain factor. Therefore, there were many “negative” results. Some of these results could be false negatives, where a factor was present but not detected. One potential source of false negatives is a lack of informative trials for an individual factor. For example, easy trials on the ends of the psychometric curve tend to be informative about the presence of guessing; thus, having too few easy trials may have prevented us from detecting guessing. Similarly, a narrow orientation range may have prevented us from detecting the oblique effect. A second potential source is trade-offs between parameters. For example, a nonzero guessing rate can be mimicked by a zero guessing rate and a lower (mean) precision parameter. To illustrate this trade-off, we generated a synthetic data set with the G model for Experiment 4, with a precision of 0.08 deg^−2^ and a guessing rate of 0.02, and computed the log likelihood with different combinations of precision and guessing rate in the G model. Different combinations of precision and guessing rate fit the data equally well, including a precision with 0 guessing rate (**Figure 9A**). In such a scenario, the LL_max_ of the with-factor model (G) could be identical to the LL_max_ of the without-factor model (Base) even though the factor is present. In another example, in Experiments 9-11, V might trade off against O and/or D, and the weaker evidence for factor V might be due to stronger evidence for factors O (9 and 10) or D (Experiment 11). To illustrate this scenario, we generated a synthetic data set with the V model for Experiment 9, with a scale parameter *τ* = 0.05, and computed the log likelihood of different combinations of *τ* and *β* of the OV model. A combination of zero *β* and the true *τ* fit as well as different combinations of a non-zero *β* and a smaller *τ* (**Figure 9B**). A smaller fitted *τ* indicated weaker evidence for factor V, because the data were partly explained by the factor O. Trade-offs can happen with any model comparison metric, but AIC and BIC are specifically known to be insensitive to trade-offs between factors (Gelman, Hwang, & Vehtari, 2014).

**Figure 9.**
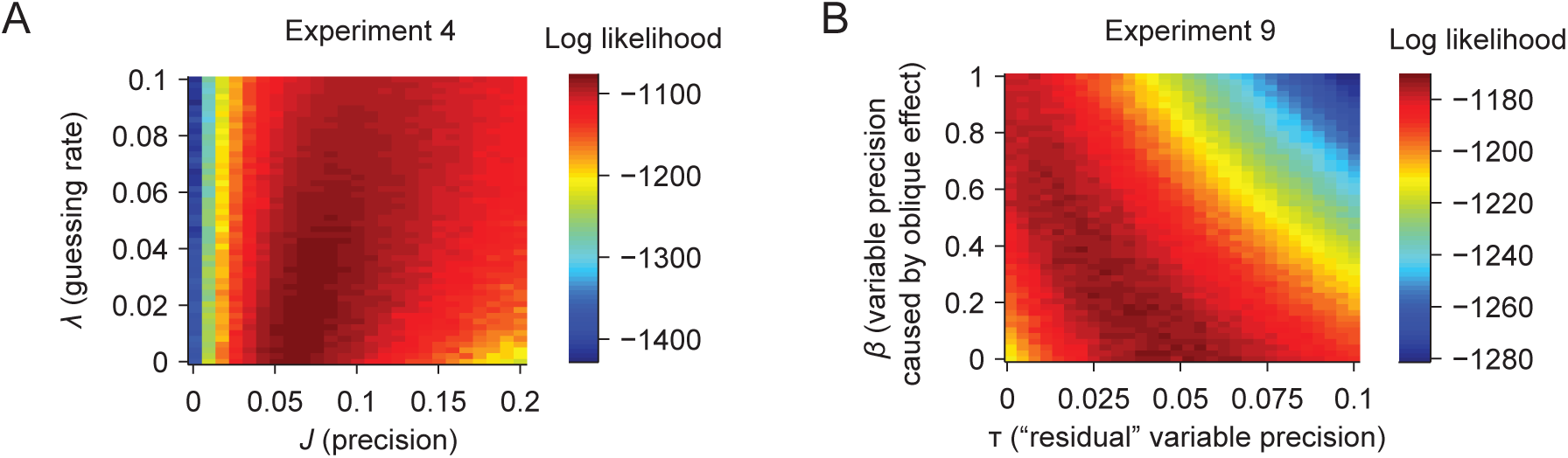
Trade-offs between parameters. **(A)** Trade-off between precision *J* and guessing rate ***X***. We generated a synthetic data set from the G model in Experiment 4 with *J*=0.08 deg^−2^ and λ*=0.02*, and fitted the data with the G model. The color plot shows the log likelihood of combinations of *J* and ***X***. Many combinations have a high log likelihood, including a combination of λ*=0* and a value of *J* lower than the true value. **(B)** Trade-off between the factors O (parametrized by β) and V (parametrized by τ). We generated a synthetic data set from the V model in Experiment 9, with 7=0.05 (and β=0), and fitted the data with the OV model. The color plot shows the log marginal likelihood of combinations of β and τ. Many combinations have a high log likelihood, including a combination of non-zero β and a value of τ lower than the true value.

### Suboptimality

We performed factorial analysis on four factors and ended up with 16 models in total. However, there were still a large number of models we did not cover. For example, a subject might have used a suboptimal decision rule in performing some of the tasks. An example of a suboptimal rule would be the max rule (Baldassi & Verghese, 2002; Eckstein, 1998; Green & Swets, 1966; Nolte, 1967; Palmer, 1990), in which the subject performs the task only based on the stimulus with the largest tilt. Another case arises when subjects have incorrectly or incompletely learned the class-conditioned stimulus distributions in the experiment, *p*(*s*|*C*=-1) and *p*(*s*|*C*=1), yet perform Bayesian inference under those wrong beliefs. We address both issues in this section.

#### Combining the GODV family with suboptimal decision rules in Experiment 7, 9 and 10

In the paper where Experiment 7 was originally presented (Shen & Ma, 2016), we systematically tested the optimal decision rule and 24 suboptimal decision rules; however, of the four factors G, O, D, and V, we only included factor G. In the present paper, we tested all factors, but so far assumed an optimal decision rule. Considering both the suboptimal rules and models in the “GODV family” simultaneously would potentially change conclusions in both studies. First, if a “bad” suboptimal rule in the previous study fits the data well now by combining with a model in the GODV family, it would imply a major change in the previous paper. Second, if a model without factor V in the GODV family fits the data well now by combining with a suboptimal rule, it would imply a change in the evidence for factor V in the current paper.

To explore these, we crossed the suboptimal rules from Shen & Ma (2016) with all members of the GODV family in this current study, when the crossing was valid. This led to 292 extra models (**Figure 10A**, for a more detailed description, see **Appendix 2**). The results confirmed the conclusions from Shen & Ma, (2016) that human behaviors are closer to optimality than to simplicity in this task (**Figure 10A**, **Figure A12**): regardless of which GODV family member the decision rule was crossed with, a) simple rules (Class I and Class II) fitted the data worse than the optimal decision rules, with mean AIC or BIC differences greater than 40; b) the more similar a suboptimal rule was to the optimal rule, the better it fitted the data (**Figure 10A**).

**Figure 10.**
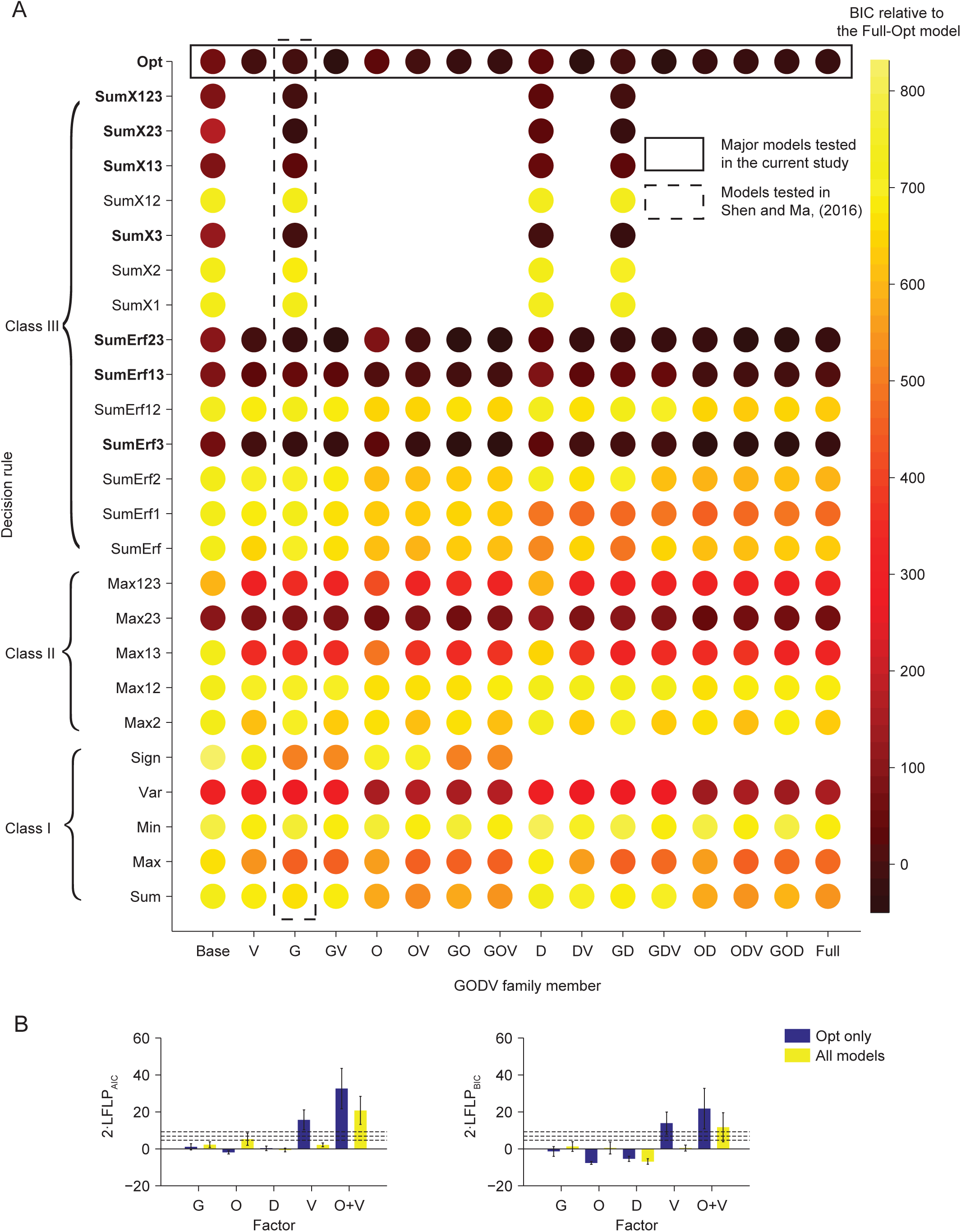
Crossing the suboptimal decision rules with the factor models in Experiment 7. **(A)** The x-axis lists GODV family members, and the y-axis lists different decision rules from Shen & Ma, 2016. The color of the dot represents the BIC of a hybrid model with a certain decision rule and factor model. Some combinations are missing because those models are invalid (Appendix 2). **(B)** Mean and s.e.m of 2·LFLR based on AIC (left) or BIC (right) for factors G, O, D, V, and the OV combination in Experiment 7. Blue bars: only models with the optimal decision rule are included. Yellow bars: all models except for those crossed with the Sign rule and SumX rules are included; we marginalized over decision rule in the same way as we marginalized over the “missing” GODV factors.

**Figure A12.**
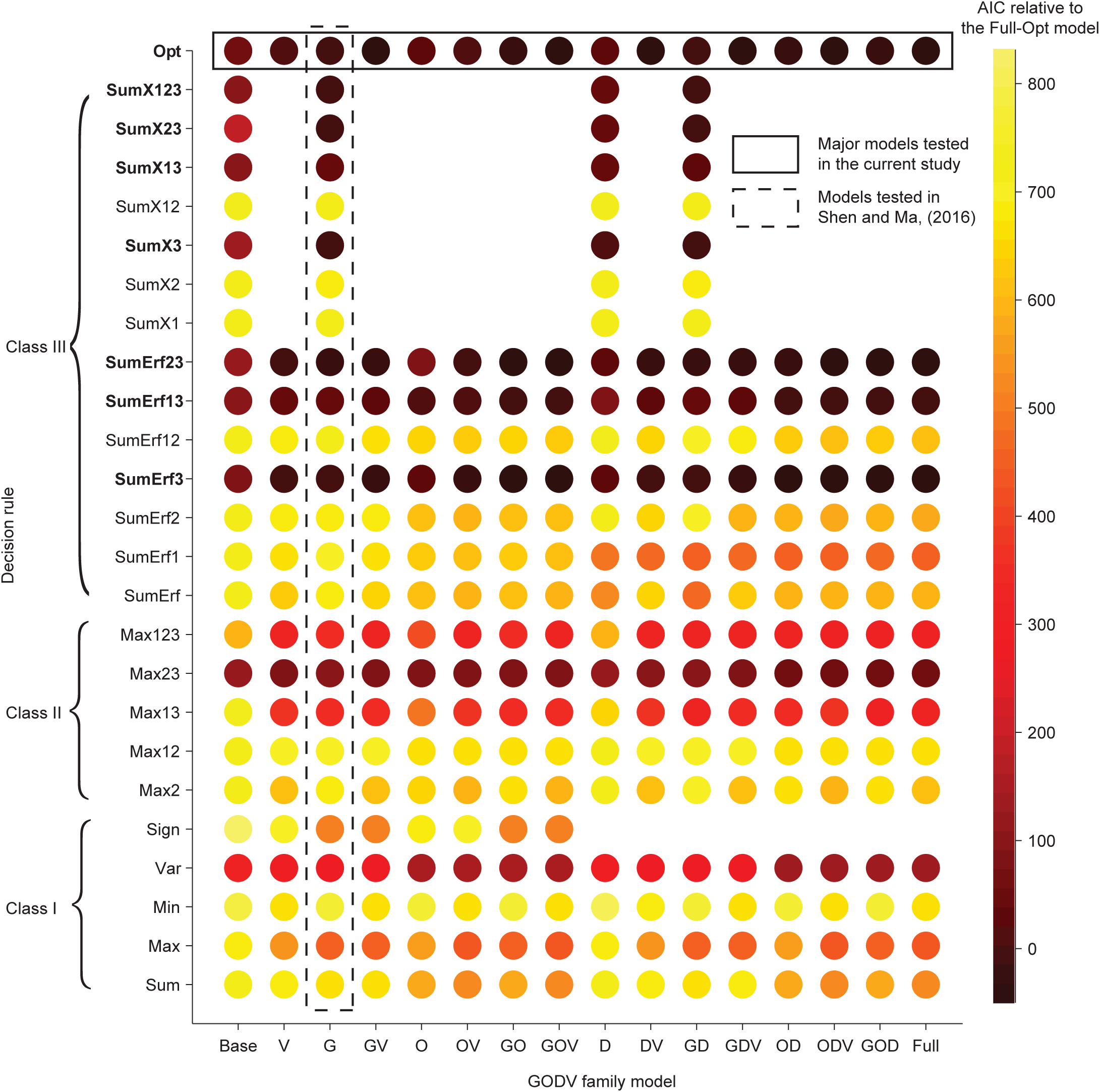
Crossing the suboptimal decision rules with the GODV factor models in Experiment 7. As Figure 10A, but computed with AIC. Results are similar to those with BIC.

By contrast, considering the suboptimal decision rules in Experiment 7 changed the earlier conclusion of the current paper about the presence of factor V. We computed 2·LFLR_AIC_(V) and 2·LFLR_BIC_(V) by marginalizing over all models, including those with a suboptimal decision rule (**Figure 10B**). The evidence for the presence of factor V decreased to 2.1 ± 1.1 (AIC) and 0.3 ± 1.6 (BIC). Thus, we hardly found any evidence for the presence of residual variable precision in any of our experiments.

In Experiments 9 and 10, the distribution of the target orientation was narrower than that of each distractor orientation. Therefore, an intuitive alternative to the optimal decision rule is to report the tilt of the least tilted stimulus. Following Shen & Ma, (2016), we call this rule the “Min” rule. This rule is not just intuitive but can also be considered a Bayesian “two-step” rule: first pick the target by maximizing *p*(*L*|**x**), where *L* is the hypothesized target location, and then report the tilt at the best location 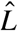, which is equivalent to maximizing *p*(*C*|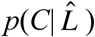). We tested combinations between the Min rule and all models in the GODV family in both Experiment 9 and 10. We found that the Min rule fitted much worse than the optimal rule regardless of the GODV family member it was crossed with (**Figure 11A, C**). This result suggests that subjects do not use the Min rule in these two experiments. Consistently, considering both optimal rule and Min rule did not change the evidence for the presence of factors in Experiments 9 and 10 (**Figure 11B, D**).

**Figure 11.**
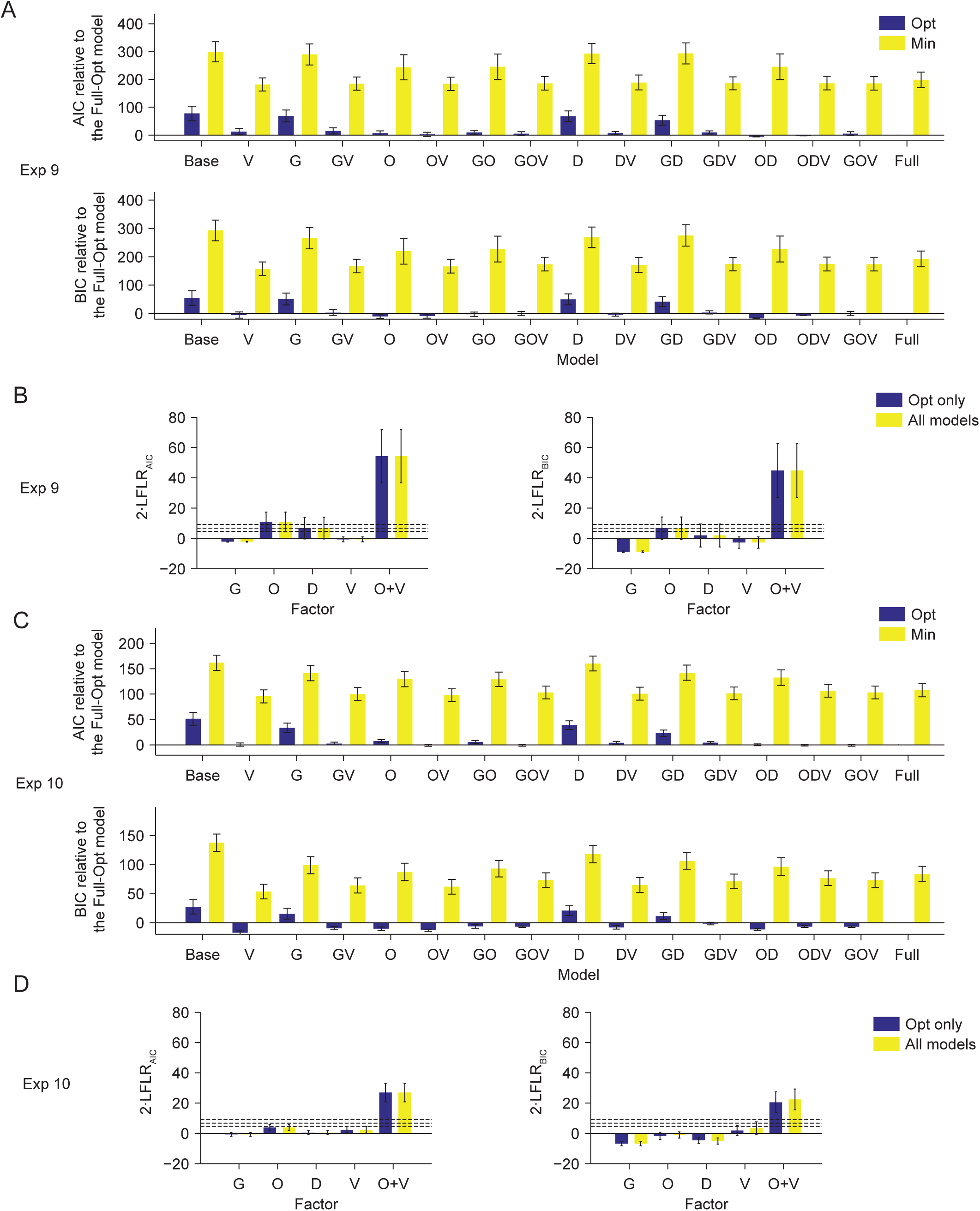
Comparing the optimal with the Min rule in Experiments 9 and 10. **(A)** Mean and s.e.m. of the difference in AIC (top) and BIC (bottom) between each model and the Full-Opt model in Experiment 9. Blue bar: models with the optimal decision rule. Yellow bar: models with the Min decision rule. **(B)** Mean and s.e.m of 2·LFLR based on AIC (left) or BIC (right) for the factors G, O, D, V, and the OV combination in Experiment 9. Blue bars: only models with the optimal decision rule are included. Yellow bars: all models are included; we marginalized over decision rule (Opt/Min) in the same way as we marginalized over the “missing” GODV factors. **(C-D)** As (A-B), but for Experiment 10.

#### Models with boxcar prior in Experiment 5

In Experiments 1 and 3-6, where the orientation of the target was discrete, we did not model the prior over target orientation as the true target orientation distribution, because it is unlikely that subjects learned a dense, discrete distribution. Instead, we modeled the prior to be a zero-mean Gaussian distribution with the same standard deviation of the true distribution, which was a form of suboptimality. Alternatively, we could assume a boxcar prior of the target distribution that covered the orientation range of the true distribution. We tested all GODV family members with this prior and compared the results to the GODV family members with the Gaussian prior in Experiment 5. We found that both AIC and BIC were similar between the Gaussian prior and the boxcar prior (**Figure 12A**), and changing the prior did not change the evidence for the factors (**Figure 12B**). This result suggests that our conclusions are not sensitive to what prior we use when the target orientation is discrete.

**Figure 12.**
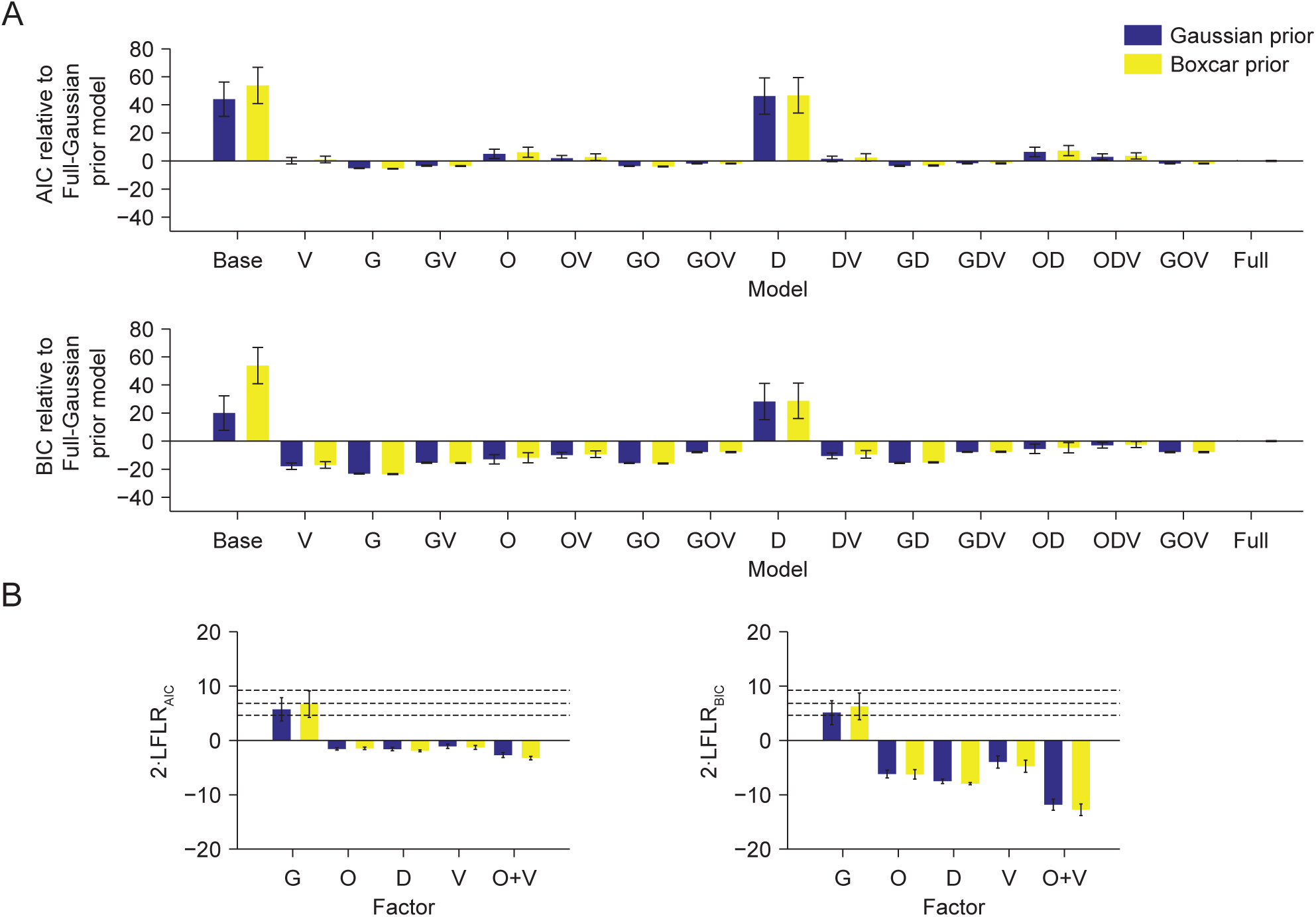
Comparing models with a Gaussian prior and a boxcar prior over orientation in Experiment 5. **(A**) Mean and s.e.m. of the difference in AIC (top) and BIC (bottom) between each model and the Full-Gaussian prior model in Experiment 5. Blue bars: models with a Gaussian prior. Yellow bar: models with a boxcar prior. **(B)** Mean and s.e.m of 2·LFLR based on AIC (left) or BIC (right) for the factors G, O, D, V, and the OV combination in Experiment 5. Blue bar: models with a Gaussian prior. Yellow bars: models with a boxcar prior.

**Figure 13.**
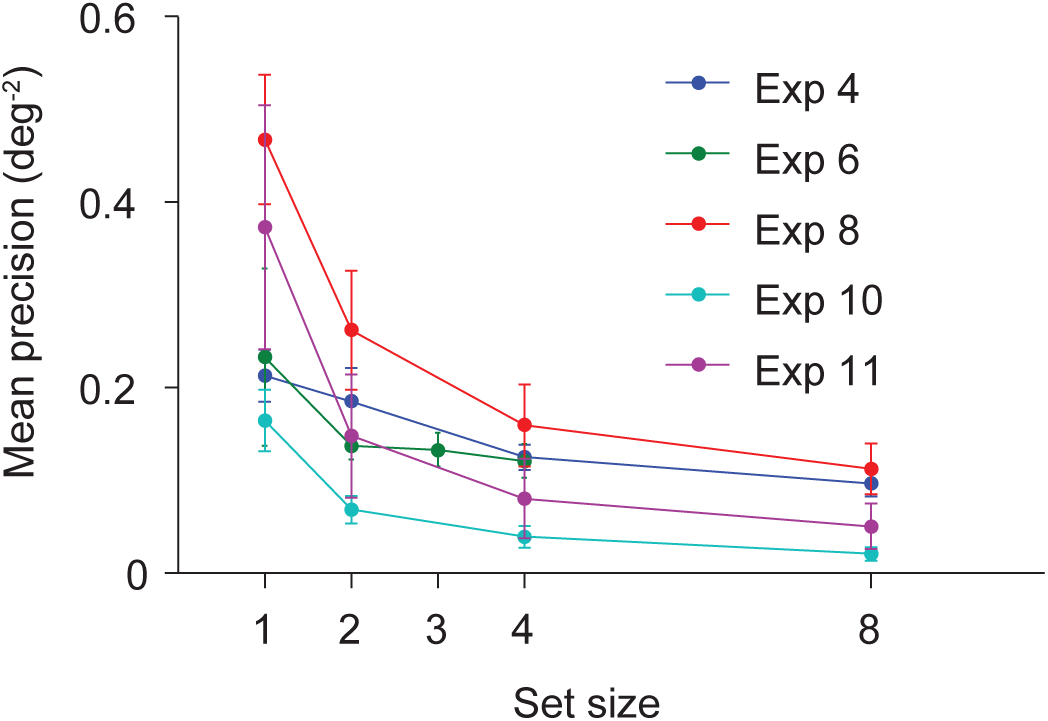
Relationship between mean precision and set size, estimated with the Full model, in all experiments with multiple set sizes (mean ± 1 s.e.m.). The effect of set size is significant in all experiments except Experiment 6.

### Relationship between mean precision and set size

Experiments 4, 6, 8, 10, and 11 used multiple set sizes, allowing us to explore the effects of task on the relationship between mean precision and set size. Mean precision, as estimated in the Full model, decreased strongly with set size in Experiments 8, 10 and 11 (repeated-measures ANOVAs: *p* < 0.05); in these experiments, the distractors were variable across trials. There were no obvious differences between detection (Experiments 8 and 11) and categorization (Experiment 10). There was no significant effect of set size in Experiment 6 (*F*(3, 6) = 1.1, *p* = 0.38), where the distractors were fixed at vertical (**Figure 13**). In Experiment 4, all stimuli were targets but with an orientation that was unpredictable across trials. Although performance increased with set size (**Figure A13A**; *F*(3, 6)=7.25, *p* < 0.01), because more stimuli gave more information about the correct answer, we found that mean precision *decreased* with set size (**Figure 13**: *F*(3, 6) = 4.18, *p* = 0.013). Given the weak evidence found for factor G in Experiment 4, we also estimated the precision with the ODV model, but the set size effect was similar (**Figure A13B**: *F*(3, 6) = 6.07, *p* < 0.01).

**Figure A13.**
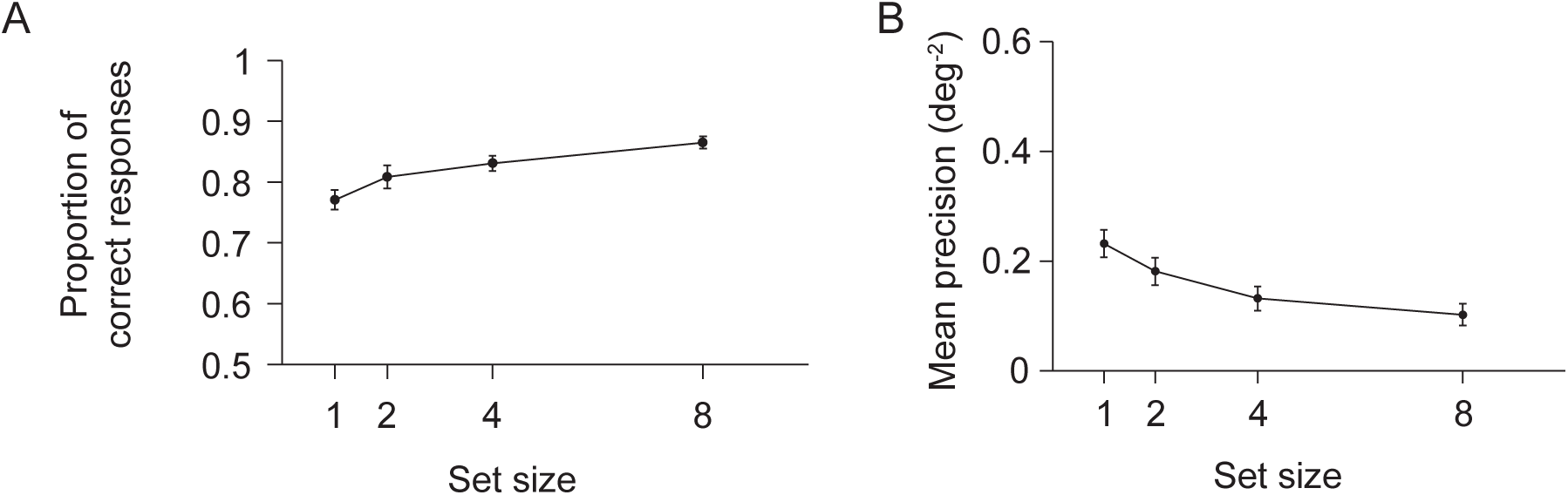
Effects of set size in Experiment 4. Even though proportion correct increases as a function of set size (A), mean precision *decreases* with set size both when estimated with the Full model (Fig. 13) and with the ODV model (B). Error bars denote ± 1 s.e.m. <u>Click here to access/download</u>

Experiments 8 and 11 were from Mazyar et al. (2013) (Experiment 2 and Experiment 1, respectively), and even though there were minor differences between the models, the relationship between mean precision and set size was very similar as in the original paper. An earlier paper (Mazyar et al., 2012) considered one more visual search condition. When the distractors were fixed at 5°, mean precision was constant across different set sizes. Based on the results of both studies, the latter paper hypothesized that mean precision decreases with set size if the *distractors* are unpredictable across trials. The results from Experiments 6 and 10 are broadly consistent with this conclusion. However, the design of Experiment 4 was not covered by this hypothesis: there were no distractors but yet we found a significant effect of set size. A unifying hypothesis could be that the less predictable the *entire stimulus display* is across trials, the stronger the decrease of mean precision with set size. However, in all of this, one needs to keep in mind the possibility that the estimates of the precision parameters are affected by tradeoffs with guessing (**Figure 9A**).

Ultimately, it would be more satisfactory to have a normative explanation: *why* does mean precision decrease with set size to different extents for different stimulus statistics? One recent proposal is that set size effects are due to an optimal trade-off between behavioral performance and the neural costs associated with stimulus encoding (van den Berg & Ma, 2017). Greater predictability might allow for more efficient neural coding, which would lead to savings in neural cost, and that in turn to a weaker set size effect.

## DISCUSSION

### Summary

We asked whether variable precision exists in visual perception. Specifically, we varied the complexity of the distractor context. We analyzed data from 11 visual experiments that used very similar oriented stimuli, and performed factorial model comparison with six factor importance metrics. Overall, we found little evidence for residual variable precision (V) when accounting for guessing (G), the oblique effect (O), and decision noise (D). In Experiments 7 through 11, if we had only considered factors G and V, we would have claimed evidence for the presence of factor V. However, when we considered factors O and D as well, the evidence weakened in Experiment 8 and disappeared in Experiments 9 through 11 (consistent with findings by Pratte et al., (2017) that were obtained without factorial model comparison). Evidence for the presence of factor V remained strong in Experiment 7, but then disappeared when considering suboptimal decision rules. Thus, we are not convinced that precision is ever variable in visual perception. On the positive side, this means that modelers of visual perception might not making a major mistake when they not include variable precision in their models.

### Why did we not get stronger results?

#### Group evidence

We quantified all evidence for models or factors by taking the average of AIC, BIC, or derived metrics over subjects. Instead, we could have summed (K. E. Stephan, Marshall, Penny, Friston, & Fink, 2007). We deliberately did not do so, because the underlying assumption would have been that all subjects follow the same model, which is something that we are not convinced of. A solution could have been to do group Bayesian model selection (Rigoux, Stephan, Friston, & Daunizeau, 2014; klaas Enno Stephan, Penny, Daunizeau, Moran, & Friston, 2009), which marginalizes over assignments of subjects to models. However, we did not trust this method given the low numbers of subjects and large numbers of models in our experiments. Therefore, we decided to describe evidence in an even more conservative way, namely by averaging over subjects. This approach has led us to conclusions that are certainly more cautious than if we had summed evidence, and probably more cautious than if we had used group Bayesian model selection.

#### Experimental design

We did not optimize our stimulus design for our main question of whether residual variable precision is present. For example, we could have calculated the target or distractor distribution width that would have had the highest expected information gain about the presence of factor V, by simulating large numbers of synthetic data sets. However, it is not clear whether this would have helped a lot, and moreover, it would have depended on the assumptions of subject parameters. In addition, asking subjects to report their confidence could increase model identifiability (van den Berg et al., 2017).

### Relation to previous work

#### Relation to work on visual short-term memory (VSTM)

Recent studies that claimed to find evidence for variable precision in VSTM (D. T. Devkar et al., 2015; Fougnie et al., 2012; keshvari et al., 2012, 2013; van den Berg et al., 2012) did not take into account all confounding factors that we considered here: guessing, heteroskedasticity and decision noise. This raises the question of how much residual variability is present in VSTM when accounting for all confounding factors. One study has suggested that oblique effect accounts for most of what otherwise would have been designated variable precision (Pratte et al., 2017), but that there is still evidence for the presence of residual variable precision; however, decision noise was not considered. If evidence for the presence of residual variable precision would survive consideration of all other factors, it would suggest that the residual variable precision is memory-related.

In color short-term memory, estimation precision was found to be much higher for some color configurations than for others, beyond what would be expected from heteroskedasticity (Brady & Alvarez, 2015). This raises the possibility that stimulus context is critical for residual variable precision. In the present work however, we did not find evidence for factor V even when the stimulus configuration was very different across trials (in Experiments 7-11). One possible explanation for this discrepancy is that delayed estimation might be more sensitive to the presence of residual variable precision than our binary categorization task. Unfortunately, delayed estimation is not easily adapted to a purely perceptual setting. Another possibility is that residual variability in precision is greater in working memory than in perception.

#### Relation to work on discriminating noise in different stages

Previous work has characterized noise in human behavior with various approaches. In contrast detection studies, varying external noise allows one to estimate internal noise (Burgess et al., 1981; Liu et al., 1995; Pelli & Farell, 1999). In Burgess et al., (1981), Pelli & Farell, (1999), this method is based on a linear relationship between threshold signal energy and noise energy. They then define the intercept to be the “internal noise” and the slope to be the “sampling efficiency”. The “internal noise” roughly corresponds to sensory noise in our framework, although the noise in the decision stage would also contribute. “Sampling efficiency” characterizes how close to optimal the decoder is, e.g. how well matched Gabor filters are to the stimulus1. Like other forms of suboptimality, low sampling efficiency could cause more variability in behavior; in our models, it would be absorbed into decision noise (Beck et al., 2012).

More recently, Drugowitsch et al. (2016) distinguished sources of suboptimality in an evidence accumulation task. The factors they tested included encoding noise, inference noise, selection noise, and deterministic biases. Their encoding noise was equivalent to ours (without O or V). Their inference noise and selection noise were both forms of decision noise, with the former being added at each time step and the latter only once at the end; in our work, these are indistinguishable. They compared models that each had one form of noise with the Base model without noise, similar to our knock-in analysis. They found that a model with inference noise explains the data best. However, they did not do full factorial model comparison and did not compute evidence of factor presence; therefore, their results cannot immediately be compared with ours.

### Factor importance in factorial model comparison

Apart from our scientific question, some of the model comparison methods we used might be useful in other contexts. Although, factorial model comparison (Acerbi, Vijayakumar, et al., 2014; van den Berg et al., 2014) helps avoid biases and oversights when deciding which models to compare, its drawbacks include model proliferation and model non-identifiability. Model proliferation is the phenomenon that the number of models rises exponentially in the number of factors (van den Berg et al., 2014). For example, in Experiment 7, we tested a total of 308 models. This large number of models makes it challenging to sensibly summarize the conclusions of the model comparison. Moreover, many models will be difficult to distinguish, or in other words, they will be *non-identifiable* (Lehmann & Casella, 1998, Definition 1.5.2) (Acerbi, Ma, & Vijayakumar, 2014; Shen & Ma, 2016; van den Berg et al., 2014).

Both problems might be alleviated by focusing on the evidence that a factor is important, rather than on the evidence for a specific model. van den Berg et al., (2014) summarized the results of their factorial model comparison into evidence curves for factors. Here, we introduced three new metrics of factor importance: KID, KOD, and LFLR. All three can be directly computed from the evidence for individual models. The third one is the most principled and reflects the evidence that a factor is present. However, we made several assumptions in calculating it: that the models tested are “representative”, that all models have equal prior probabilities conditioned on factor presence or absence, and that log marginal likelihood can be estimated from AIC or BIC. All these assumptions should be questioned, and the toolkit for quantifying evidence for factor importance will need to be further refined.

## ACKNOWLEDGMENTS

This work was funded by grant R01EY020958 from the National Institutes of Health. We thank Luigi Acerbi for sharing his Bayesian Adaptive Direct Search algorithm at an early stage, and for discussions. We thank Ronald van den Berg for advice and assistance during experimental design.

## Appendix 1: Decision rules

Notations:

Φ(x): standard normal cumulative function; *x_i_*: internal measurement of the *i* stimulus; *J_i_*: encoding precision of the *i* stimulus; *J_s_:* precision to generate stimulus orientations; *K_i_*: concentration parameter of Von Mises distribution; *κ*_s_: concentration parameter of the Von Mises distribution to generate stimulus orientations; *p*_right_, *p*_clockwise_, or *p*_present_: prior probability of reporting “right”, “clockwise” or “present”; *N:* set size.

In Experiments 5 to 11, where there is uncertainty about the target location, the optimal observer considers each location to be a possible location of the target, and weighs these possibilities (hypotheses) by their probabilities:

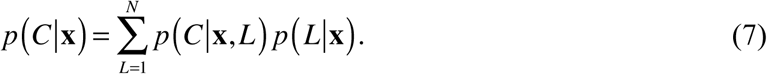

*Experiment 1:*

The observer reports “right” when

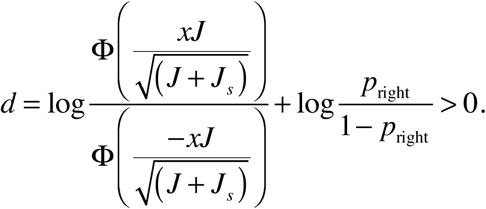

*Experiment 2:*

The observer reports “clockwise” when

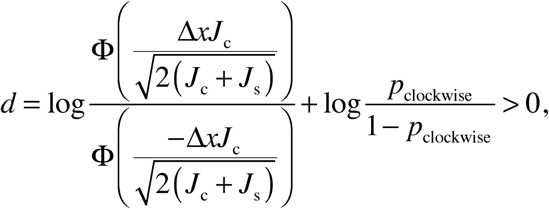

where 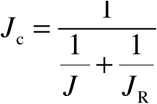. *J*_R_ denotes the encoding precision of the reference, and *J* denotes the encoding precision of the target. ∆*x* denotes the internal measurement of the target relative to that of the reference.

*Experiments 3 and 4*

The observer reports “right” when

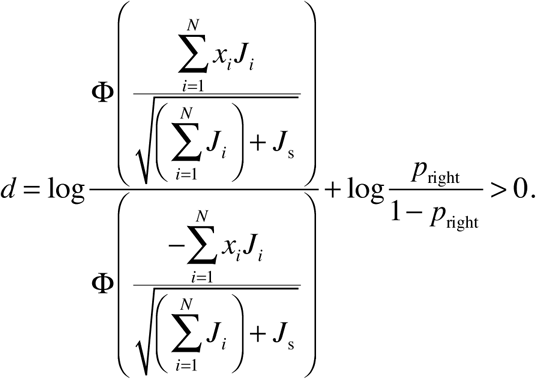

*Experiments 5 and 6*

The observer reports “right” when

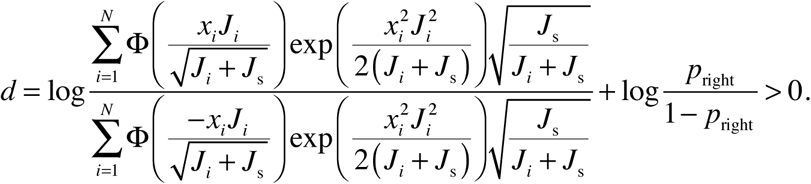

*Experiment 7*

The observer reports “right” when

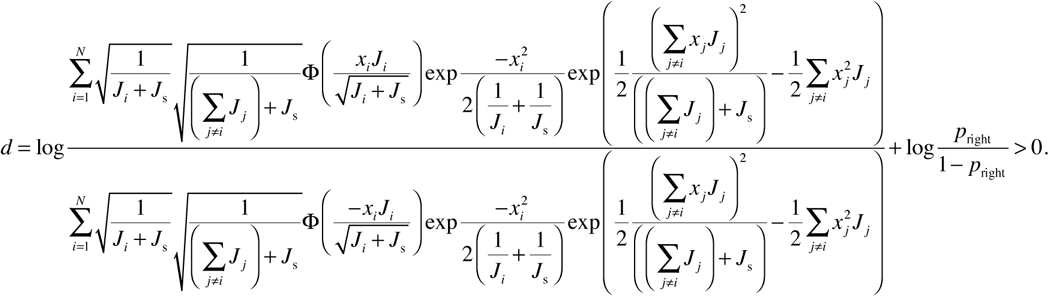

*Experiment 8*

The subject reports “present” when

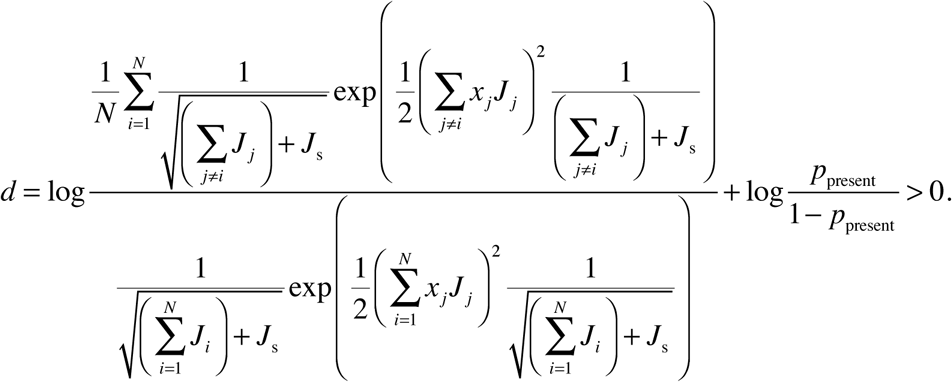

*Experiments 9 and 10*

The subject reports “right” when

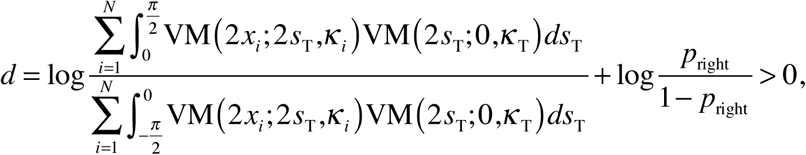

where VM(*x*; *s, κ*) denotes a Von Mises distribution with a mean of *s* and concentration parameter of *κ, s*_T_ denotes the target orientation, and *κ*_T_ denotes the concentration parameter of the Von Mises distribution to generate target orientations.

*Experiment 11*

The subject reports “present” when

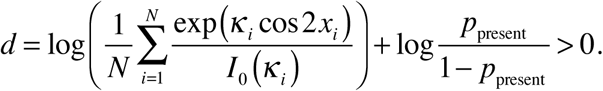

## Appendix 2: Combinations of the GODV family with suboptimal rules in Experiment 7

In Shen & Ma, (2016) where Experiment 7 was first published, we tested three classes of suboptimal rules: Class I contained “simple” suboptimal rules, Class II contained “two-step” rules in which the observer first decides on target location and then reports the tilt of the purported target (thereby ignoring target uncertainty), and Class III encompassed variations of the optimal rule. All decision rules took the form “report “right” when *d* > 0”, where *d* is the decision variable.

Here, we created new models by combining these suboptimal decision rules with the GODV factors. Moreover, we included in the derivation of the decision rule a prior probability that the target was tilted right in the models where this was possible (the SumErf models in Class III). To combine the suboptimal rules with factor D, we added Gaussian noise with standard deviation σ_d_ (a free parameter) to *d*. We left out several invalid combinations: a) we combined the Sign rule in Class I only with models without factor D, because *d* in the Sign model is a small integer rather than continuous; b) we combined the SumX rules in Class II only with models Base, G, D, GD, because those rules are only compatible with fixed precision.

## Footnotes

1 Confusingly, Liu et al., 1995 also measure efficiency by varying the external noise, but in their terminology, any form of inefficiency is purely a consequence of the internal noise.

